# The Kaufman oculocerebrofacial syndrome protein Ube3b regulates synapse number by ubiquitinating Ppp3cc

**DOI:** 10.1101/672923

**Authors:** Mateusz C. Ambrozkiewicz, Silvia Ripamonti, Ekaterina Borisova, Manuela Schwark, Theres Schaub, Bekir Altas, Rüstem Yilmaz, Lars Piepkorn, Stephen Horan, Olaf Jahn, Ekrem Dere, Marta Rosário, Guntram Borck, Hannelore Ehrenreich, Katrin Willig, JeongSeop Rhee, Victor Tarabykin, Hiroshi Kawabe

## Abstract

Kaufman oculocerebrofacial syndrome (KOS) is a severe autosomal recessive disorder characterized by intellectual disability, developmental delays, microcephaly and characteristic dysmorphisms. Biallelic mutations of *UBE3B*, encoding for a ubiquitin ligase E3B are causative for KOS. In this report, we characterize neuronal functions of its murine ortholog *Ube3b*. We show that Ube3b regulates dendritic branching in a cell-autonomous manner. Moreover, *Ube3b* knockout (KO) neurons exhibit increased density and aberrant morphology of dendritic spines, altered synaptic physiology and changes in hippocampal circuit activity. Dorsal forebrain-specific *Ube3b* KO animals show impaired spatial learning and alterations in social interactions. We further demonstrate that Ube3b ubiquitinates the catalytic γ-subunit of calcineurin (Ppp3cc), the overexpression of which phenocopies the loss of *Ube3b* with regard to dendritic spine density. This work provides insights into the molecular pathologies underlying intellectual disability-like phenotypes in a genetic mouse model for KOS.

## INTRODUCTION

Neuronal development comprises a series of tightly regulated morphoregulatory processes that underlie the organization of neuronal circuits. Neurons born from ventricular progenitors undergo multipolar-to-bipolar transitions, migrate radially, and distribute horizontally in the cortical plate in an inside-first-outside-last manner. Neurons extend their axons and dendrites to form the framework of neuronal networks in the cortex. At the stage of synaptogenesis, immature synapses form upon physical contacts between developing axons and dendritic spines. More than half of such synapses are then eliminated to establish the architecture of a mature neuronal network (Tønnesen and Nägerl, 2016). Processes underlying network formation involve cell adhesion proteins, extracellular ligands, their receptors on the plasma membrane, and downstream signaling cascades mediated by, among others, small GTPases, calcium, or phosphorylation. Recent accumulating evidence reveals critical roles of ubiquitination as well as ubiquitin-like modifiers in the formation of neuronal networks (Ambrozkiewicz and Kawabe, 2015; Kawabe and Brose, 2011).

Ubiquitination involves conjugation of 8.6 kDa globular ubiquitin (Ub) to a substrate protein in a cascade of enzymatic reactions catalyzed by an E1 Ub activating enzyme, an E2 Ub conjugating enzyme, and an E3 Ub ligase. Among these enzymes, E3 ligases confer the specificity of ubiquitination by interacting with their substrate proteins (Hershko et al., 1983).

Lysine (K) residues on substrate proteins can be conjugated with a single Ub moiety (monoubiquitination) or with polyUb chains (polyubiquitination). Each of seven K residues on Ub itself can serve as an acceptor for the donor Ub to form polyUb chains. Importantly, the K residue utilized for chain formation specifies the functional consequence of ubiquitination. The K48- and K63-linked polyUb chains constitute the major types of polyUb chains in the mouse and human brains (Kaiser et al., 2011). Conjugation of K48-linked chain destines substrate proteins for proteasomal degradation, whereas functional consequences of K63-linked chain conjugation include trafficking of transmembrane proteins and lysosomal degradation (Hicke and Riezman, 1996).

E3 Ub ligases fall into two major families based on their domain architecture: Really Interesting New Gene (RING)-type and homologous to E6-AP C-terminus (HECT)-type ligase families. RING-type ligases form a transient trimeric protein complex with the E2-Ub conjugate and its substrate to facilitate transfer of Ub from E2 to the substrate acting as a scaffold protein. HECT-type ligases covalently conjugate a Ub moiety to a cysteine residue in their C-terminal catalytic domain, and thereupon transfer Ub onto their substrate proteins. While RING-type enzymes form the preponderant E3 ligase family composed of some hundreds of genes, only 28 of HECT-type E3 ligases have been identified in the human genome (Huibregtse et al., 1995).

We have previously reported that HECT-type ligases control fundamental neuronal morphoregulatory processes, such as neuronal polarity, migration, and neuritogenesis (Ambrozkiewicz et al., 2018; Hsia et al., 2014; Kawabe et al., 2010). Disturbances in Ub signaling have emerged as a molecular basis of numerous human neurodevelopmental disorders (Ambrozkiewicz and Kawabe, 2015). Recently, biallelic mutations in the human *UBE3B* gene, encoding for Ub ligase E3B, have been described causative for autosomal recessive Kaufman oculocerebrofacial syndrome (KOS), a distinct intellectual disability (ID) syndrome (OMIM #244450) (Basel-Vanagaite et al., 2012, 2014; Flex et al., 2013; Pedurupillay et al., 2015).

Although the involvement of UBE3B in the etiology of human diseases has been reported repeatedly, both molecular and cellular mechanisms underlying the ID in KOS still remain largely uncharted. In this report, we generated and characterized a conventional and brain-specific conditional knockout (cKO) mouse lines of the murine *UBE3B* orthologue *Ube3b*. We demonstrate that loss of *Ube3b* leads to impaired dendrite arborization and increased density of dendritic spines in CA1 excitatory neurons, and alters the electrophysiological properties of neurons and neuronal networks in the hippocampus. *Ube3b* cKO mice exhibit behavioral phenotypes, including a decline in spatial memory and superior social memory. We provide evidence that Ube3b controls dendritic spine number in hippocampal neurons by ubiquitinating one of the regulatory subunits of calcineurin, Ppp3cc.

## RESULTS, FIGURES, SUPPLEMENTARY FIGURES, AND FIGURE LEGENDS

### Loss of *Ube3b* in mice leads to developmental delay

In order to study the role of murine Ube3b, an ortholog of human UBE3B, we employed Cre-loxP technology to genetically knockout *Ube3b* in mice (Fig. S1A to S1D). In order to conventionally delete *Ube3b*, we crossed *Ube3b*^f/f^ mice with E2A-Cre driver mice. Conventional *Ube3b* knockout animals (*Ube3b*^-/-^) were born at Mendelian ratio, but were smaller than littermate controls, displayed prominent malformations of the snout and the orbital arch, sometimes preventing the animals from opening the eye, and pronounced kyphosis (Fig. S1E). *Ube3b*^-/-^ mice also presented with a shuffling gait when moving. Examination of *Ube3b*^-/-^ brains revealed an overall smaller organ as compared to the controls (Fig. S1F). *Ube3b*^-/-^ mice generated in this study died at approximately 3 weeks, likely due to frail teeth, malnutrition and/or muscle weakness. Because of these overt phenotypes in *Ube3b*^-/-^ also reported elsewhere very recently (Cheon et al., 2019), we decided to study the role of Ube3b in neurons using *Ube3b*^f/f^ mice and mice with *Emx1-*Cre^+/-^-driven conditional knockout of *Ube3b*, described below. This approach circumvents the syndromic phenotype of *Ube3b*^-/-^, which partially limits the identification of brain-specific and neuron-specific phenotypes, and prevent the analysis of mature brains. Using an antibody against the HECT domain of Ube3b we showed complete loss of Ube3b in the dorsal forebrains of *Ube3b*^-/-^ mice by Western blotting (Fig. S1G). Heterozygous animals (middle lane, Fig. S1G) displayed approximately 30 % of Ube3b level in the brain, but were phenotypically indistinguishable from the *Ube3b*^+/+^ mice.

### Ube3b is expressed in the brain and enriched at postsynaptic densities (PSDs)

We first studied the expression of the ligase in the murine brain (Fig. 1). We designed a DIG-probe recognizing an mRNA sequence spanning the middle region and a portion of HECT domain of Ube3b and performed *in situ* hybridization using sagittal brain sections from mice at embryonic day 16 (E16) and postnatal day 20 (P20) (Fig. 1A). We detected a strong hybridization signal in the ventricular zone and ganglionic eminences at E16. At P20, Ube3b mRNA was expressed throughout the entire CNS, including the cerebral cortex and hippocampus. We also studied the expression of Ube3b in each brain region by Western blotting. Ube3b protein was widely expressed throughout the brain, with prominent expression in the cerebral cortex, the olfactory bulb, the hippocampus, and the cerebellum as compared to other regions (Fig. 1B). Developmental expression profiling by Western blotting using cortical lysates revealed that Ube3b was expressed at constant low levels throughout embryonic development and its expression increased 6-fold from P0 to P21 to then reach the plateau phase (Fig. 1C, Table S1A, “Exact values, n numbers and statistics for the experiments in this report”). Such an expression pattern resembles postsynaptic proteins such as PSD-95 and Neuroligin-1 (Song et al., 1999). Next, we studied subcellular distribution of Ube3b in biochemically fractionated cerebral cortices from 6 – 7 weeks old mice (Fig. 1D). Ube3b was predominantly found in the PSD fraction, indicating its association with synaptic specialization in neurons.

**Figure 1.**
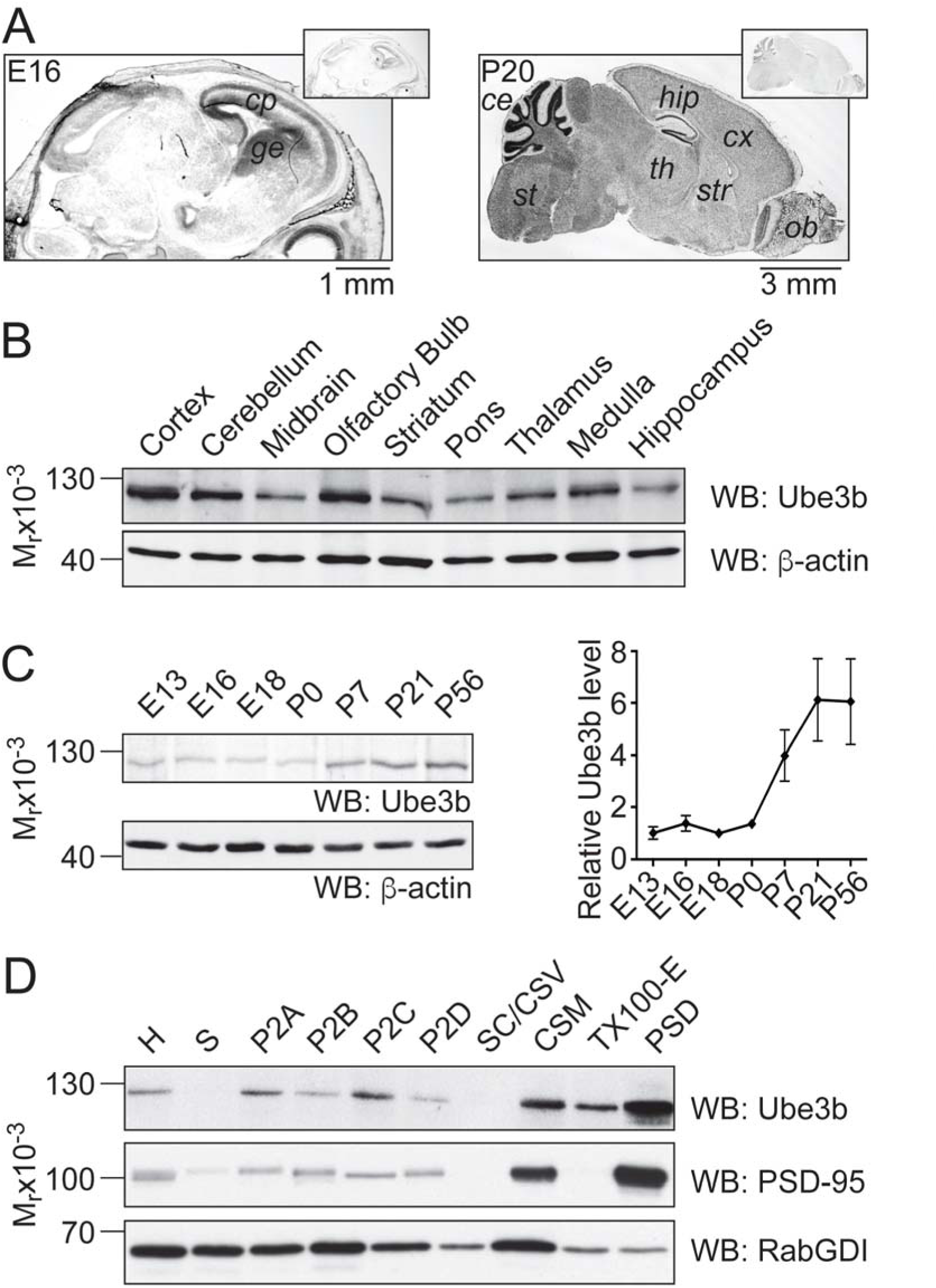
Ube3b is expressed in developing and mature murine central nervous system and associates with postsynaptic densities. (A) *In situ* hybridization on sagittal sections of E16 and P20 of mouse brains with antisense probe for mRNA of HECT domain of Ube3b (large images) and sense control probe (smaller insets). Note the enrichment of *Ube3b* mRNA in developing cortex (cortical plate, *cp*), and ganglionic eminence (*ge*), and overall abundance in the adult central nervous system (cerebellum, *ce*; brain stem, *st*; hippocampus, *hip*; cortex, *cx*; thalamus, *th*; striatum, *str*; olfactory bulb *ob*). (B) Western blotting using homogenates of adult mouse brain regions with indicated antibodies. (C) Left: developmental expression profile of Ube3b. Cortical homogenates at different developmental stages were used for Western blotting with indicated antibodies. Right: relative Ube3b level in mouse cortex during brain development. Ube3b level was normalized to β-actin and expressed relative to the ligase level in E13 embryonic cortex. Each point represents mean level from three cortices ± S.E.M (Table S1A). (D) Results of Western blotting using subcellular fractions with indicated antibodies (homogenate, H; supernatant, S; synaptic cytosol, SC; myelin fraction, P2A; ER/Golgi fraction, P2B; synaptosomes, P2C; mitochondria-enriched fraction, P2D; crude synaptic vesicles, CSV; crude synaptic membrane, CSM; TritonX100 extract, TX100-E; postsynaptic density, PSD).

Taken together, Ube3b is expressed in developing and mature pallium, where it might be an important regulator of the synapse.

### *Ube3b* knockout abrogates neurite branching in primary hippocampal neurons

Impaired dendritogenesis has been described as a cellular pathology for ID syndromes (Wang et al., 2013). In order to study the impact of *Ube3b* loss on dendrite development, we analyzed the morphology of primary hippocampal neurons prepared from *Ube3b*^f/f^ mice expressing EGFP (control), or EGFP and Cre recombinase (*Ube3b* KO). *Ube3b* KO neurons display pronounced deficits in neurite branching at fourth day *in vitro* (DIV4; Fig. 2A). Next, we infected *Ube3b*^f/f^ primary hippocampal neurons with lentiviruses expressing EGFP, or Cre recombinase and concluded that at DIV5, *Ube3b*^f/f^ neurons expressing Cre recombinase are devoid of Ube3b (Fig. 2B). At this stage, axons of *Ube3b* KO neurons developed significantly less primary branches as compared to control (Fig. 2C, Table S1B). At DIV7 (Fig. 2D), we quantified the complexity of neurites by Sholl analysis. *Ube3b* KO neurons displayed overall less complex neurite branching as compared to control cells (Fig. 2E and 2F, Table S1C).

**Figure 2.**
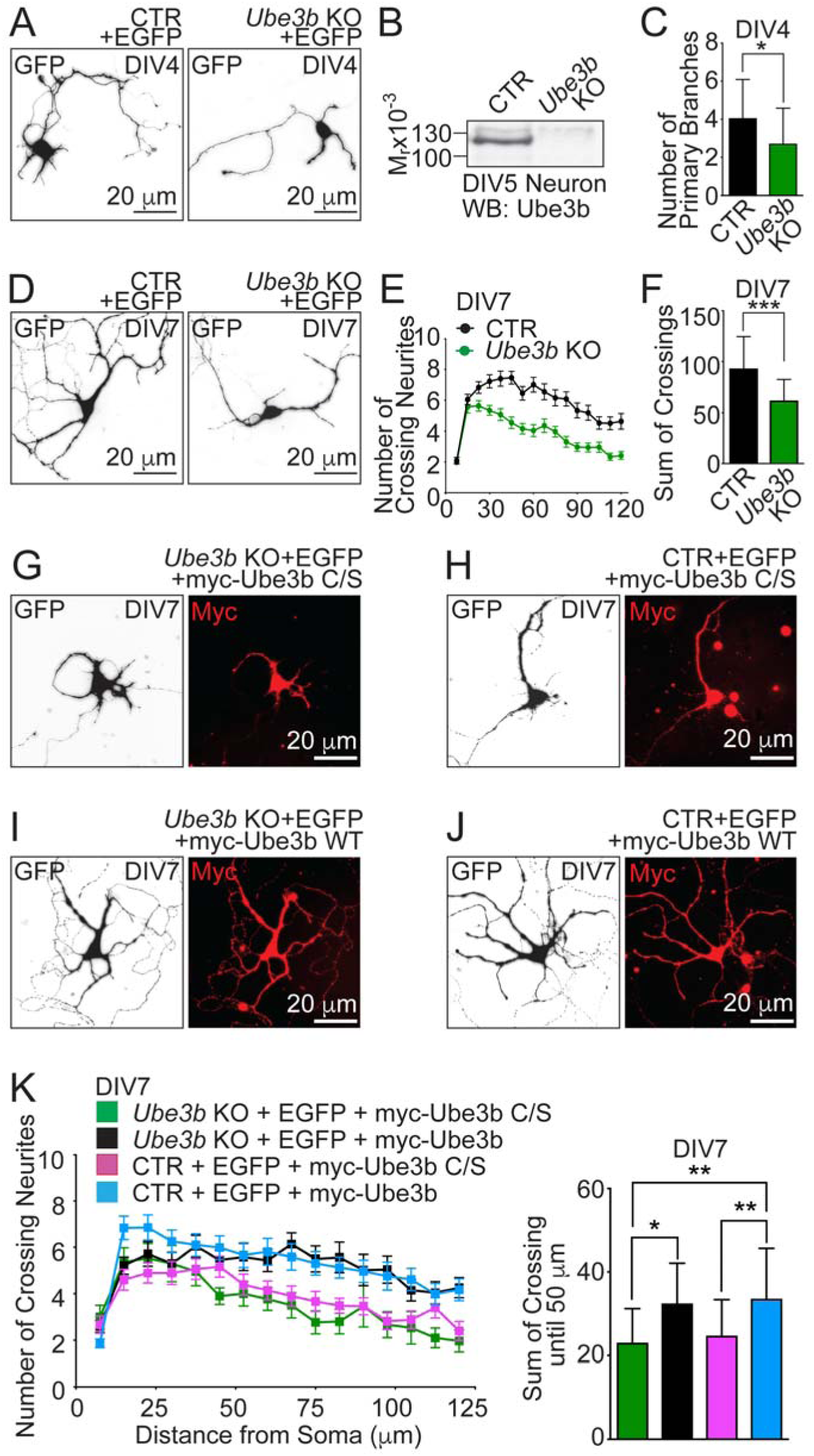
*Ube3b* loss in neurons leads to dramatic impairment of neurite branching. (A) Images of primary hippocampal neurons at day 4 *in vitro* (DIV4) prepared from *Ube3b*^f/f^ mice and expressing EGFP (control, CTR), or co-expressing EGFP and Cre recombinase to induce *Ube3b* knockout (KO), after immunostating with anti-GFP antibody. (B) Western blotting against Ube3b in neuronal lysates. Primary hippocampal neurons from *Ube3b*^f/f^ mice were infected at DIV1 with lentiviruses expressing EGFP (CTR), or co-expressing EGFP and Cre recombinase (*Ube3b* KO), and harvested at DIV5. (C) Quantification of primary axonal branches in DIV4 control and KO neurons. Axon was defined as the longest neurite (Table S1B). (D and E) Representative images (D) and Sholl analysis diagram (E) of CTR and *Ube3b* KO neurons immunostained with an anti-GFP antibody at DIV7. (F) Quantification of the sum of total crossings of neurites with Sholl circles from a single neuron (Table S1C). (G-J) Representative images of GFP and myc immunostaining from CTR or *Ube3b* KO neuron expressing EGFP and myc-tagged wild type Ube3b (myc-Ube3b) or myc-tagged catalytically inactive mutant of Ube3b (myc-Ube3b C/S). (K) Left: Sholl analysis diagram of CTR and *Ube3b* KO neurons expressing either myc-Ube3b, or myc-Ube3b C/S. Right: Quantification of the sum of total crossings of neurites with Sholl circles (Table S1D). Results on (E) and (K, left) are represented as averages ± S.E.M. Results on (C), (F), (K, right) are represented as averages ± S.D. For statistics on (C) and (F), unpaired t-test; for (K, right), one-way ANOVA with Tukey’s multiple comparisons test; *** p < 0.001; ** 0.001 < p < 0.01; * 0.01 < p < 0.05.

To study if impaired dendritogenesis in *Ube3b* KO neurons is a consequence of the loss of its enzymatic activity, we transfected *Ube3b*^f/f^ primary hippocampal neurons with plasmids encoding EGFP alone (control), or EGFP and Cre (*Ube3b* KO) together with either myc-tagged wild type Ube3b (Fig. 2I, and 2J), or with the catalytically inactive point mutant of Ube3b (Fig. 2G, and 2H). In this mutant, a cysteine residue in the catalytic HECT domain acting as a Ub acceptor is substituted by serine (Ube3b C1038S, hereafter called Ube3b C/S). The expression of recombinant wild type Ube3b in *Ube3b* KO neurons restored neurite branching (compare Fig. 2D, left panel, and Fig. 2I; note that experiments shown in Fig. 2D to 2K were performed simultaneously). The expression of Ube3b C/S, on the other hand, had almost no impact on neurite complexity in *Ube3b* KO neurons (compare Fig. 2D, right panel and Fig. 2G; Fig. 2G, and 2K, Table S1D). The expression of wild type Ube3b in the control neurons had almost no effect on neurite arborization (compare Fig. 2D, left panel, and 2J), possibly because of saturating endogenous Ube3b expression level required for proper dendritic branching. Expression of Ube3b C/S in control neurons decreased the complexity of neurites (Fig. 2F, and 2K, Table S1D), indicating dominant-negative effects of Ube3b C/S overexpression.

Altogether, we demonstrate that the enzymatic activity of Ube3b is critical to promote neurite arborization.

### Dorsal telencephalon-specific *Ube3b* knockout mice display reduction of cortical thickness

In order to circumvent secondary effects caused by multitude of non-neuronal defects in *Ube3b*^-/-^ mice and to study the role of Ube3b in the forebrain, we established a glutamatergic neuron- and glia-specific conditional *Ube3b* KO (*Ube3b*^f/f^; *Emx1*-Cre^+/-^, hereafter referred to as *Ube3b* cKO) mouse line (Guo et al., 2000). In this mouse line Cre is active in glutamatergic neurons and locally born glial cells, but not in GABAergic interneurons. We confirmed significant reduction of Ube3b protein level in the cortical lysate from *Ube3b* cKO mice by Western blotting (Fig. 3A). Residual Ube3b expression comes likely from non-*Emx1* expressing cells, e.g. GABAergic inhibitory neurons.

**Figure 3.**
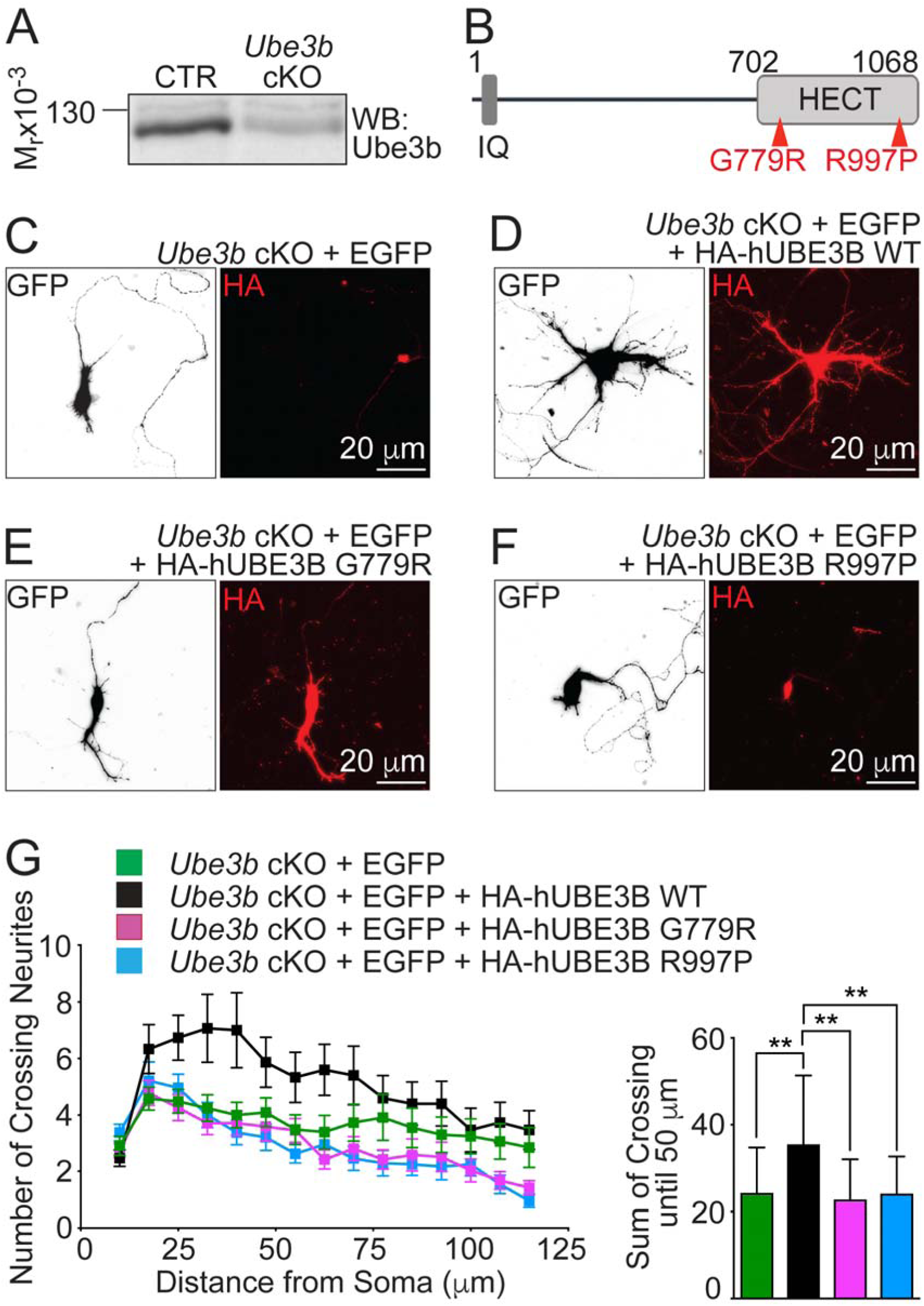
Pathogenic missense mutations in *UBE3B* identified in KOS patients abrogate neurite branching. (A) Validation of Ube3b loss in the *Emx1*-Cre^+/-^ driven conditional knockout of *Ube3b* (*Ube3b*^f/f^; *Emx1-*Cre^+/-^*, Ube3b* cKO) cortex. Western blotting using cortical lysates of adult control (*Ube3b*^f/f^) and *Ube3b* cKO mice with the antibody against Ube3b. (B) A schematic of a topography of G779R and R997P missense mutations in human UBE3B. (C to F) Primary hippocampal neurons from *Ube3b* cKO mice expressing EGFP and variants of HA-tagged human UBE3B at DIV7, after immunostaining with anti-GFP antibody. Representative images of GFP and HA staining in *Ube3b* cKO neuron expressing EGFP alone (C), EGFP with HA-tagged wild type human UBE3B (HA-hUBE3B) (D), EGFP with HA-tagged human UBE3B G779R missense mutant (HA-hUBE3B G779R) (E), EGFP with HA-tagged human UBE3B R997P missense mutant (HA-hUBE3B R997P) (F). (G) Left: Sholl analysis diagrams of *Ube3b* cKO neurons expressing indicated proteins. Right: quantifications of the sum of total crossings of neurites with Sholl circles until 50 μm from the center of the soma (Table S1E). Results on (G, left) are represented as averages ± S.E.M. Results on (G, right) are represented as averages ± S.D. For statistics on (G, right), one-way ANOVA with Tukey’s multiple comparisons test; ** 0.001 < p < 0.01.

Next, we examined the gross morphology of the brain in *Ube3b* cKO mice. Brains from 9 weeks old *Ube3b* cKO male mice reveal a reduction in cortical thickness as compared to control *Ube3b*^f/f^ mice (Fig. S2A, and S2B, Table S1F), while numbers of cortical neurons in the coronal cross sections (Fig S2C, Table S1G) and density of neurons (Fig. S2D, Table S1H) show no difference between control and *Ube3b* cKO. The reduction of the cortical thickness can therefore be attributed to the decreased dendritic branching. In some of analyzed *Ube3b* cKO brains, we noted ventricular dilatation upon *Ube3b* cKO that were not associated with increased inflammatory responses in the brain (data not shown). At P0, we detected no changes in cortical layering patters (Fig. S3), as investigated by immunostaining with upper layer markers Special AT-Rich binding protein 2 (Satb2), Cut-Like 1 (Cux1), and deeper layer marker Chicken ovoalbumin upstream promoter transcription factor-interacting protein 2 (CTIP2; Fig. S3A and S3B). Furthermore, immunostaining for NCAM-L1, a marker for callosally projecting axons showed intact projection fibers and unchanged thickness of the *corpus callosum* at the midline in *Ube3b* cKO mice (Fig. S3C). The discrepancy from KOS patients with hypoplastic *corpus callosum* is likely due to involvement of cells of non-*Emx1* lineage in the formation of this tract in our cKO model.

In addition, we at times observed that several analyzed animals exhibited a thinner cortex at P0. This observation was always associated with enlarged lateral ventricles, likely involving hydrocephaly. However, given the intact cortical neuron number in the absence of *Ube3b*, enlarged ventricles do not seem to affect neurogenesis or neuronal survival in *Ube3b* cKO mice (Fig. S2).

### Pathogenic missense mutations in UBE3B are of loss-of-function type

Along with the truncating mutations leading to a frame shift and premature stop codon in *UBE3B*, we have previously reported two additional missense mutations in the HECT domain, Arg997Pro (R997P) and Gly779Arg (G779R) present in patients with KOS (Basel-Vanagaite et al., 2014) (Fig. 3B).

We then studied the consequences of G779R and R997P mutations for the role of Ube3b in neurite branching. We prepared primary hippocampal neurons from *Ube3b* cKO mice and expressed EGFP as a control (Fig. 3C), or co-expressed EGFP and HA-tagged wild type human UBE3B (HA-hUBE3B; Fig. 3D), EGFP and HA-tagged G779R mutant (HA-hUBE3B G779R; Fig. 3E), or EGFP and HA-tagged R997P mutant of hUBE3B (HA-hUBE3B R997P; Fig. 3F). Overexpression of human UBE3B in *Ube3b* cKO neurons restored neurite branching as compared to EGFP expressing *Ube3b* cKO neurons at DIV7 (Fig. 3C, 3D, and 3G). Strikingly, expression of either missense mutants of UBE3B failed to rescue defective neurite branching (Fig. 3C-3G, Table S1E) at this stage. Of these mutants, HA-hUBE3B R997P displayed a peculiar perinuclear localization (Fig. 3F). In conclusion, UBE3B residues G779 and R997, mutated in patients with KOS, are essential for its function in neurite development and R997 is crucial for proper subcellular distribution of UBE3B in the nerve cell.

### *Ube3b* KO leads to increased density and altered spine morphology in hippocampal neurons

One of the common features characteristic for ID syndromes in humans, such as Angelman syndrome and in ID-mouse models is altered spine number and/or spine morphology (Dindot et al., 2008). We next studied the density and morphology of dendritic spines in CA1 hippocampal neurons in control and *Ube3b* cKO mice. We transfected hippocampal progenitors *in utero* at E14.5 with EGFP- and myristoylated Venus-encoding plasmids to visualize dendritic spines. Myristoylated Venus labels plasma membranes of transfected cells, providing an advantageous way of visualizing dendritic spines for morphometry. Brains were fixed at P21, when synaptogenesis in the mouse brain has been accomplished and dendritic branches stabilized (Koleske, 2013).

We noted a dramatic increase in spine density on primary branches of apical dendrites in *Ube3b* cKO neurons as compared to control, using confocal microscopy (Fig. 4A). In order to perform spine morphometrics with super-resolution microscopy we employed stimulated emission depletion (STED) microscopy (Fig. 4B-4H). We detected a marked increase in spine density in *Ube3b* cKO neurons as compared to control (Fig. 4B and 4D, Table S1I). In order to test if this is due to cell autonomous role of Ube3b, we conducted morphometric analysis of spines in Ube3b-deficient neurons surrounded by cells with wild type levels of Ube3b. In these experiments, we deleted *Ube3b* by transfecting small number of progenitors in *Ube3b*^f/f^ with low concentration of Cre expression plasmid (*Ube3b* KO). Similarly, *Ube3b* KO led to an increase in spine density (Fig. 4C and 4E, Table S1J). No differences concerning the frequency of spine types between control and *Ube3b* KO neurons were observed (Fig. 4F, Table S1K). Necks of spines in *Ube3b* KO neurons were longer (Fig. 4C and 4G, Table S1L), and spine heads wider as compared to control (Fig. 4C and 4H, Table S1M). In the further analyses of Ube3b in formation and maintenance of dendritic spines, we focused on spine density changes, a phenotype consistent in *Ube3b* cKO and *Ube3b* KO and the most robust among all morphological alterations.

**Figure 4.**
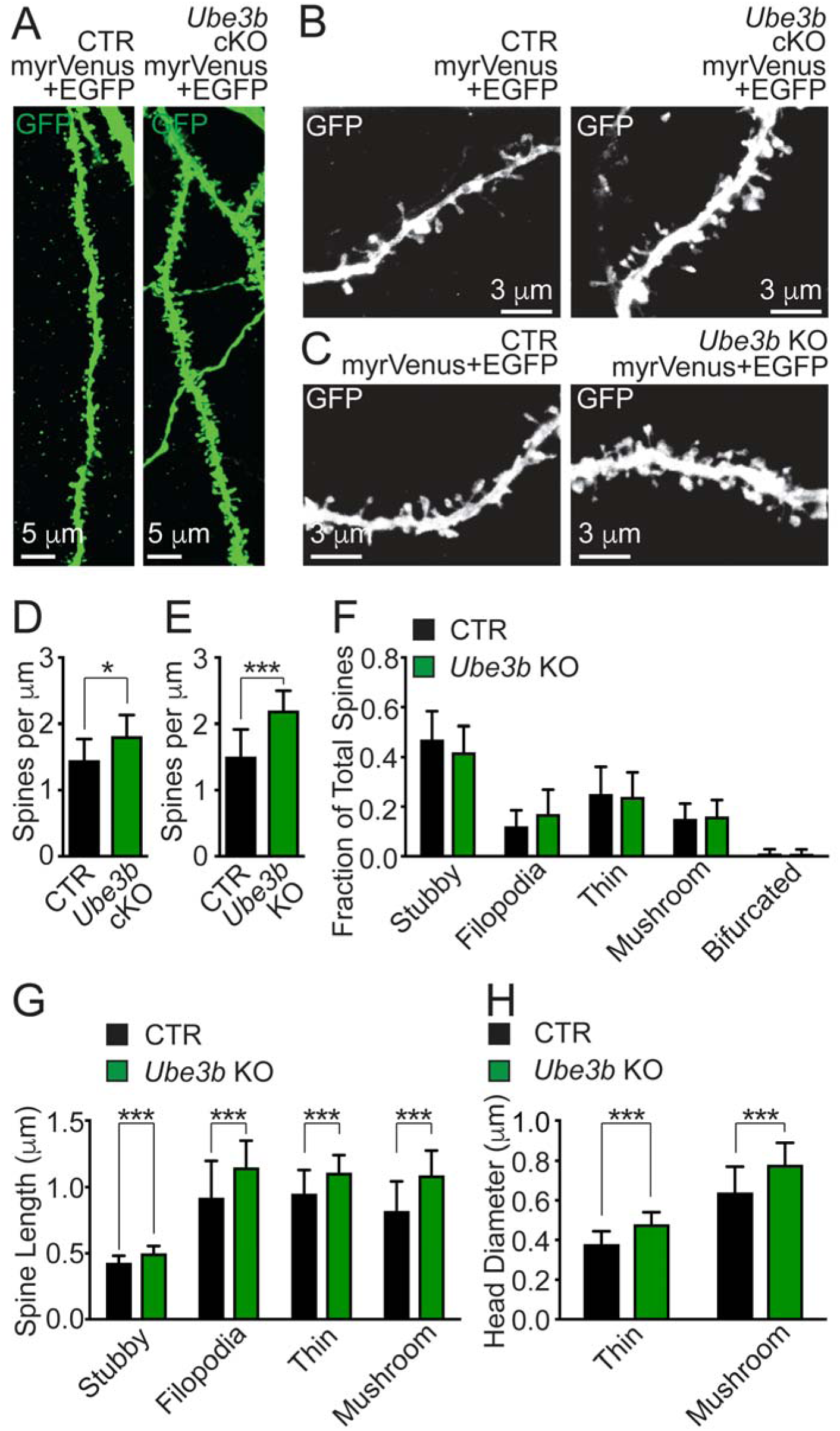
*Ube3b* deletion in neurons induces morphological aberrances in dendritic spines *in vivo*. (A) Representative images of anti-GFP immunostaining signal from primary apical branches of CA1 pyramidal neurons in *Ube3b*^f/f^ (CTR) and *Ube3b*^f/f^; *Emx1*-Cre^+/-^ (*Ube3b* cKO) mice at P21. Hippocampal progenitors were *in utero* electroporated at E14.5 with plasmids encoding for EGFP and myr-Venus. Myr-Venus labels neuronal membranes, allowing for visualization of dendritic spines. (B, C) Maximum intensity projections of STED images of anti-GFP immunostaining signal from dendrites of CA1 pyramidal neurons of CTR (left panel) and *Ube3b* cKO (right panel) mice (B), and *Ube3b*^f/f^ neurons expressing EGFP and myr-Venus (CTR; left panel) or EGFP, myr-Venus, and Cre recombinase (*Ube3b* KO; right panel) (C). (D, E) Quantification of spine densities in CTR and *Ube3b* cKO neurons (Table S1I) and in CTR and *Ube3b* KO neurons (Table S1J). (F) Frequencies of morphological groups of spines in CTR and *Ube3b* KO neurons (Table S1K). (G, H) Quantification of spine length measured from the emergence of spine on the dendrite until the head tip (G; Table S1L) and head diameter of thin, and mushroom spines (H; Table S1M). Results on bar graphs are represented as averages ± S.D. For statistics on (D, E) and (G, H), unpaired t-test; (F), two-way ANOVA with with Bonferroni post-hoc test; *** p < 0.001; * 0.01 < p < 0.05.

We also analyzed *Ube3b* KO neurons at P10, when dendritic spines are initially specified on developing neurons. Expression of myc-tagged Ube3b in *Ube3b* KO neurons rescued spine density, indicative of a cell-autonomous role of the ligase in regulating spine numbers (Fig. S4, Table S1O). Altogether, Ube3b emerges as a pivotal Ub ligase controlling spine number and morphology in hippocampal neurons.

### *Ube3b* KO leads to an increase in the number of functional excitatory synapses in hippocampal neurons

Given the branching deficiencies and spine morphology alterations, we then studied the effects of *Ube3b* loss on synaptic transmission. We analyzed glutamatergic autaptic hippocampal neurons from control and *Ube3b* cKO with electrophysiological recordings (Fig. 5) (Bekkers and Stevens, 1991).

**Figure 5.**
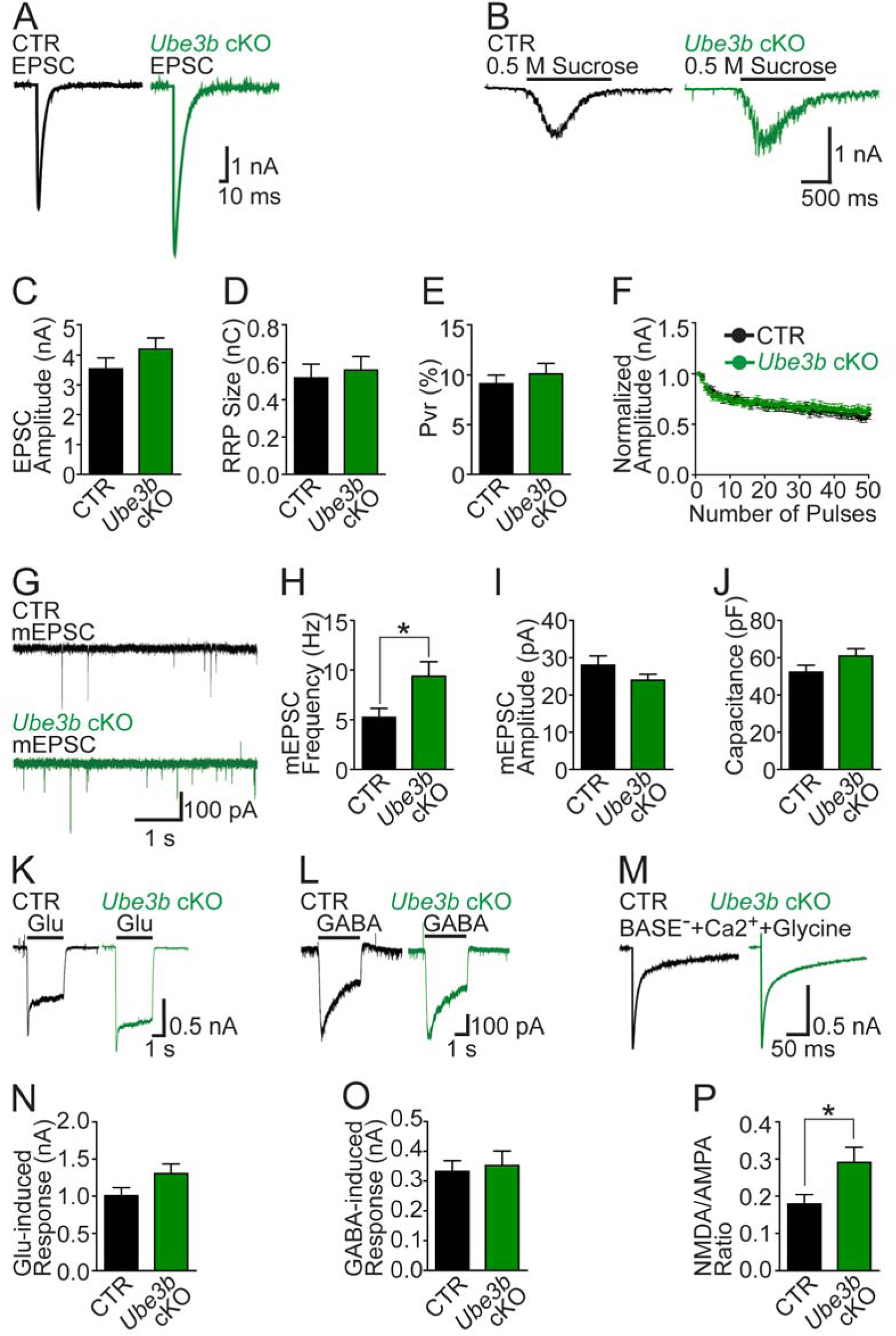
Synaptic transmission is altered in *Ube3b* cKO autaptic neurons. (A) Sample traces of evoked excitatory postsynaptic currents (EPSCs). (B) Sample traces of postsynaptic currents following hypertonic sucrose application in CTR and *Ube3b* cKO neurons. (C) Quantification of amplitude of evoked EPSCs (Table S1P). (D) Quantification of readily releasable vesicle pool (RRP) size (Table S1Q). (E) Quantification of vesicular release probability (Pvr) (Table S1R). (F) Plots of normalized peak amplitude versus number of stimuli at 10 Hz frequency. (G) Sample traces of miniature EPSCs (mEPSCs) recorded in the presence of 300 nM tetrodotoxin. (H) Quantification of mEPSCs frequency (Table S1S). (I) Quantification of mEPSCs amplitude (Table S1T). (J) Quantification of membrane capacitance (Table S1U). (K-M) Sample traces of postsynaptic currents induced by application of 100 μM exogenous glutamate (Glu) (K), of 3 μM GABA (L), and upon activation of both - NMDARs and AMPARs by Mg^2+^-free extracellular solution containing 1.8 mM CaCl_2_ and 10 μM glycine (M). (N) Quantification of glutamate induced responses (Table S1V). (O) Quantification of GABA-induced responses (Table S1W). (P) Quantification of ratio between current amplitudes through NMDARs and AMPARs (Table S1X). Results on bar graphs are represented as average ± S.E.M. For statistics, unpaired t-test; * p < 0.05.

Evoked excitatory postsynaptic currents (EPSCs, Fig. 5A) and the readily releasable pool (RRP) size (Fig. 5B) did not show significant differences in *Ube3b* cKO neurons as compared to controls (Fig. 5C and 5D, Table S1P and S1Q). Accordingly, the vesicular release probability (Pvr) (Fig. 5E, Table S1R) and short-term plasticity (Fig. 5F) were unaffected upon *Ube3b* cKO.

The frequency of miniature EPSC (mEPSC), reflecting spontaneous and action potential (AP)-independent release of the neurotransmitter (Fig. 5G), was dramatically increased (Fig. 5H, Table S1S) in *Ube3b* cKO neurons compared to control while the amplitude of mEPSCs was unaltered (Fig. 5I, Table S1T). Additionally, we detected no differences in plasma membrane capacitance, reflecting no differences in soma sizes between control and *Ube3b* cKO neurons (Fig. 5J, Table S1U). These findings illustrate that upon *Ube3b* loss, the number of functional synapses is increased, while the number of receptors on a single dendritic spine, the quantal size, and the size of the cell body are unaltered.

In order to examine the relative number of postsynaptic receptors expressed on the cell surface, we recorded postsynaptic currents induced by the bath application of glutamate (Glu) (Fig. 5K), γ-aminobutyric acid (GABA) (Fig. 5L), or Mg^2+^-free, Ca^2+^-and-glycine-containing solution (Fig. 5M) in control and cKO neurons. We detected a non-significant trend towards an increase in current amplitude following exogenous glutamate application in *Ube3b* cKO neurons (Fig. 5N, Table S1V) and no differences in the response to exogenous GABA (Fig. 5O, Table S1W). α-amino-3-hydroxy-5-methyl-4-isoxazolepropionic acid receptor (AMPAR)- and N-methyl-D-aspartate receptor (NMDAR)-mediated currents were measured in Mg^2+^-free solution (Fig. 5M). Interestingly, we observed an increase in NMDA/AMPA ratio for *Ube3b* cKO neurons as compared to control (Fig. 5P, Table S1X). Given that cell surface AMPA receptors are not changed in *Ube3b* cKO neurons (Fig. 5N), this result indicates that *Ube3b* loss leads to an increase in functional NMDA receptors expressed on the synaptic plasma membrane.

### *Ube3b* cKO mice show altered local network activity in the hippocampus

Increased density of morphologically different spines on glutamatergic neurons may alter the properties of hippocampal circuitry in *Ube3b* cKO animals, affecting the ability of neurons to synchronize in the local circuit. In order to validate this hypothesis, we recorded extracellular field potentials in the CA3 region in acute brain slices from P14-P24 animals, and induced γ-oscillatory activity by application of kainate (KA) (Fig. 6A and 6B). Notably, bath application of 100 nM KA (Ripamonti et al., 2017) resulted in epileptiform activity in approximately 85 % of slices prepared from *Ube3b* cKO mice (Fig. 6C, Table S1Y), consistent with the observation of seizures in some patients with KOS. Lowering KA to 50 nM resulted in a decrease in the number of slices with epileptic seizure-like activity and enabled recording of the γ-oscillations. While no changes in the frequency of γ-oscillations were detected between control and *Ube3b* cKO animals (Fig. 6D, Table S1Z), we observed an increase in the average power of γ-oscillations in *Ube3b* cKO as compared to control mice (Fig. 6E, Table S1A’). These data indicate that *Ube3b* cKO alters the physiology of hippocampal circuitry.

**Figure 6.**
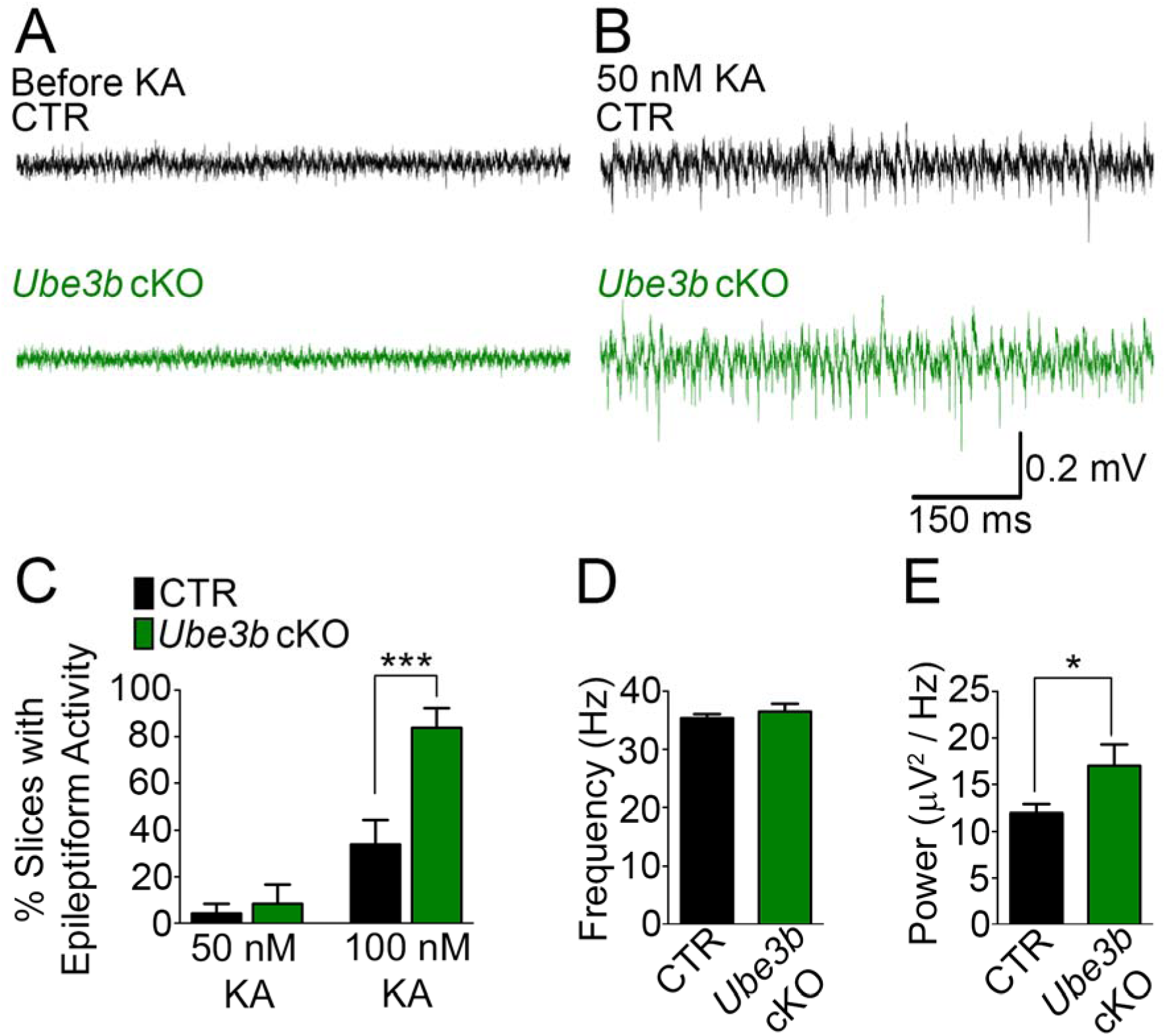
*Ube3b* cKO mice show aberrant γ-oscillations. (A and B) Representative traces of extracellular field recordings from the CA3 region of acute brain slices before (A) and after (B) the treatment with 50 nM kainate (KA) in *Ube3b*^f/f^ (CTR) and *Ube3b*^f/f^; *Emx1*-Cre^+/-^ (*Ube3b* cKO). (C) Quantification of the number of brain slices in CTR and *Ube3b* cKO mice with epileptiform activity (Table S1Y). (D and E) Quantification of the frequency (D; Table S1Z) and power (E; Table S1A’) of γ-oscillations in CTR and *Ube3b* cKO mice. Results on bars represent average ± S.E.M. For statistics, unpaired t-test; * p < 0.05; *** p < 0.001.

### *Ube3b* is critical for spatial memory, social memory and social interaction in mice

To analyze behavioral changes upon loss of *Ube3b* in the brain, we performed a battery of behavioral examinations of control (CTR; *Ube3b*^f/f^) and *Ube3b* cKO mice (Fig. 7 and Fig. S5).

**Figure 7.**
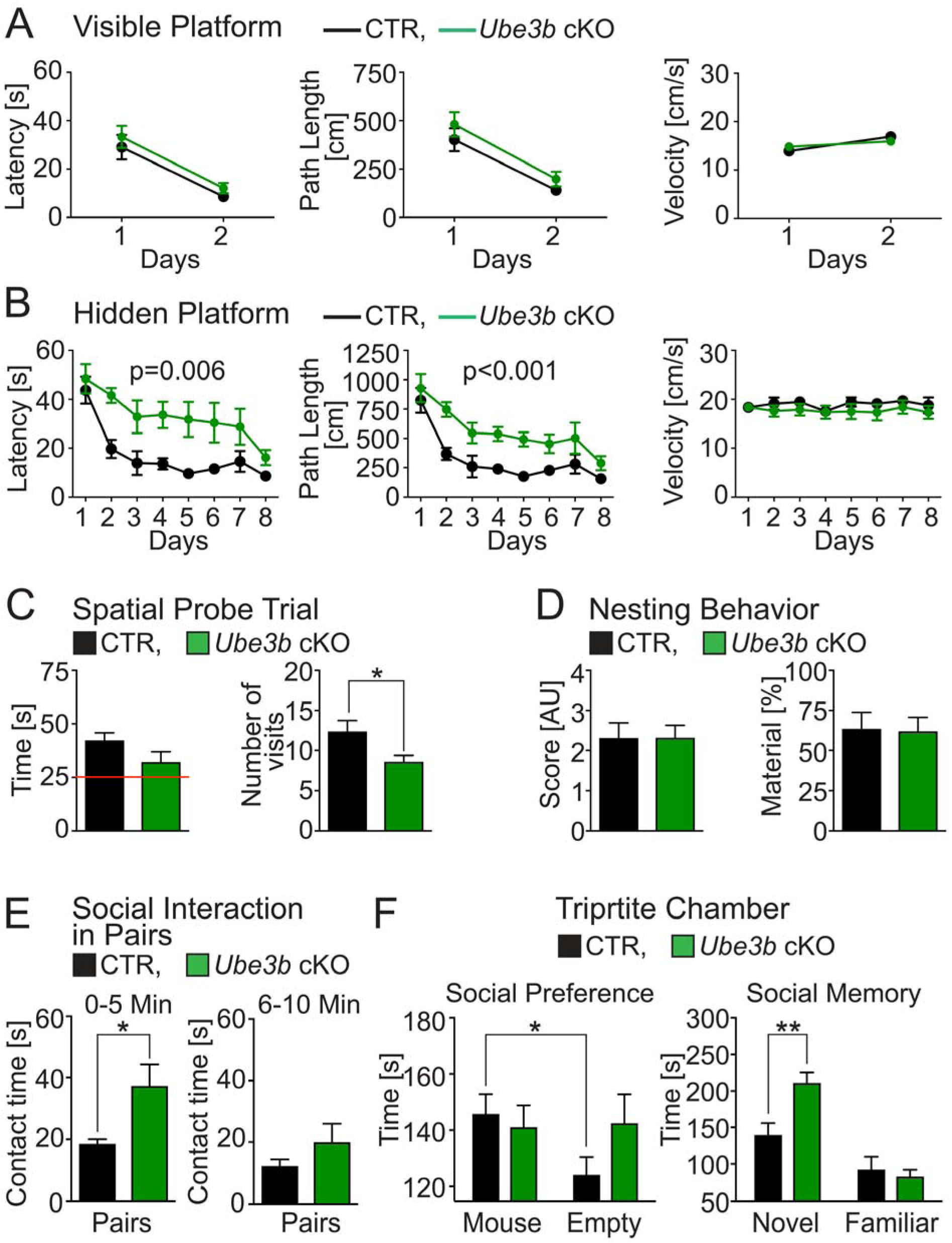
*Ube3b* cKO male mice show loss of spatial memory and increased sociability. (A) Visible platform test. Quantification of escape latencies, path length to reach the visible platform, and swim speed in CTR and *Ube3b* cKO mice (Table S1C’). (B) Morris Water Maze test with a hidden platform. Quantification of escape latencies, path length to reach the hidden platform, and swim speed in CTR and *Ube3b* cKO mice (Table S1D’). (C) Spatial Probe Trial. Quantification of the time and the number of visits of CTR and *Ube3b* cKO mice spent in target zone. Red line indicates performance at chance likelihood (Table S1E’). (D) Nesting behavior. Quantification of nesting score and the material used for building a nest in control and *Ube3b* cKO (Table S1F’). (E) Social interaction in pairs test. Quantification of contact time between two unfamiliar mice of the same genotype during the first and during the last five minutes of the test (Table S1G’). (F) Tripartite chamber tests. Quantifications of times spent with the unfamiliar mouse versus empty compartment, and with the unfamiliar versus familiar mouse (Table S1H’). All described tests were conducted with males. Results on graphs represent average ± S.E.M. For statistics on (A) and (B), Repeated Measures ANOVA; (C-E) and (F, right graph), unpaired t-test; (F, left graphs), paired t-test; ** 0.001 < p < 0.01; * 0.01 < p < 0.05.

The analysis of somatosensory functions (Fig. S5A) revealed no defects in vision or olfaction but a reduced pain threshold in *Ube3b* cKO animals. Behavioral phenotyping of freely moving animals in the home cage revealed decreased locomotion and increased frequency in self-directed behaviors, such as grooming and eating (Fig. S5B). Motor coordination tests revealed no changes in *Ube3b* cKO animals in rotarod and open field test (Fig. S5C). Consistent with the results from the home cage, during exploratory tasks, cKO animals exhibited reduced locomotion. There were no signs of increased anxiety, or impulsivity in *Ube3b* cKO animals. Pre-pulse inhibition test showed that *Ube3b* cKO animals exhibit increased processing of higher intensity stimuli as compared to control (Fig. S5D, Table S1B’).

We then subjected the mice to behavioral tests that require elaborate pallium-dependent cognitive functions. First, we investigated spatial learning measured for eight subsequent days, when mice navigated to a platform submerged underneath the water. Importantly, the visible platform test showed intact procedural learning in mice upon *Ube3b* loss (Fig. 7A, Table S1C’). In the hidden platform test, *Ube3b* cKO mice showed increased escape latencies (Fig. 7B) and swam longer distances in the pool before finding the platform than the control. Observed spatial learning deficits in *Ube3b* cKO animals were independent of changes in the motivation, motor or sensory-motor integration deficits (Table S1D’). Upon removal of the platform from the water-maze, *Ube3b* cKO mice spent less time in the area where the platform was previously located (Fig. 7C) and made fewer visits to the former platform quadrant (Table S1E’) than the control.

Next, we analyzed social interactions, where a pair of unfamiliar mice of the same genotype is placed into a familiar environment to measure their propensity to engage in social interactions. During the first five minutes of the test, *Ube3b* cKO mice spent more time together as compared to the control mice, indicative of increased sociability (Fig. 7E, Table S1G’).

In the tripartite chamber test for sociability and social memory, control mice showed a preference for the chamber with the unfamiliar mouse over the empty chamber, but no such bias was observed in the *Ube3b* cKO mice (Fig. 7F). Additionally, during the social memory test on the third trial, the *Ube3b* cKO exhibited superior social memory and spent more time in the compartment with the unfamiliar mouse as compared to the chamber with the familiar mouse (Fig. 7F, Table S1H’).

*Ube3b* cKO mice exhibit complex behavioral phenotype with both gains and losses of function, including a facilitated processing of intensive, high-arousal sensory stimuli, behavioral switch towards self-directed actions, impaired hippocampus-dependent spatial learning and memory, increased sociability, and superior social memory.

### Ppp3cc is a substrate of Ube3b Ub ligase

According to its domain structure and published data (Braganza et al., 2017), Ube3b acts as an E3 Ub ligase. Indeed, in an autoubiquitination assay, Ube3b formed Ub K48- and K63-linked polyUb chains (Fig. S6A). This indicates that Ube3b mediates proteasomal and/or lysosomal degradation. Given the enrichment of Ube3b at the PSD fraction (Fig. 1F), we hypothesized that Ube3b-mediated ubiquitination and subsequent degradation of substrates takes place locally at the synapse. To identify Ube3b substrates, we purified synaptic plasma membrane fractions (SM3) from control and *Ube3b* cKO mice (Fig S6B) to identify proteins upregulated in *Ube3b* cKO samples by comparative mass spectrometry. Based on mass spectrometric label-free quantification data, we obtained a list of synaptic proteins upregulated in SM3 fractions from *Ube3b* cKO (Fig. 8A, Table S2, “Label free quantification of proteins in SM3 fractions from control and *Ube3b* cKO cortices”). We then corroborated that the top hit protein γ-catalytic subunit of protein phosphatase 2B, Ppp3cc, was upregulated in *Ube3b* cKO synaptic membrane samples by quantitative Western blotting (Fig. 8B and 8C; Table S1I’), indicating that Ppp3cc may be a locally regulated synaptic Ube3b substrate.

**Figure 8.**
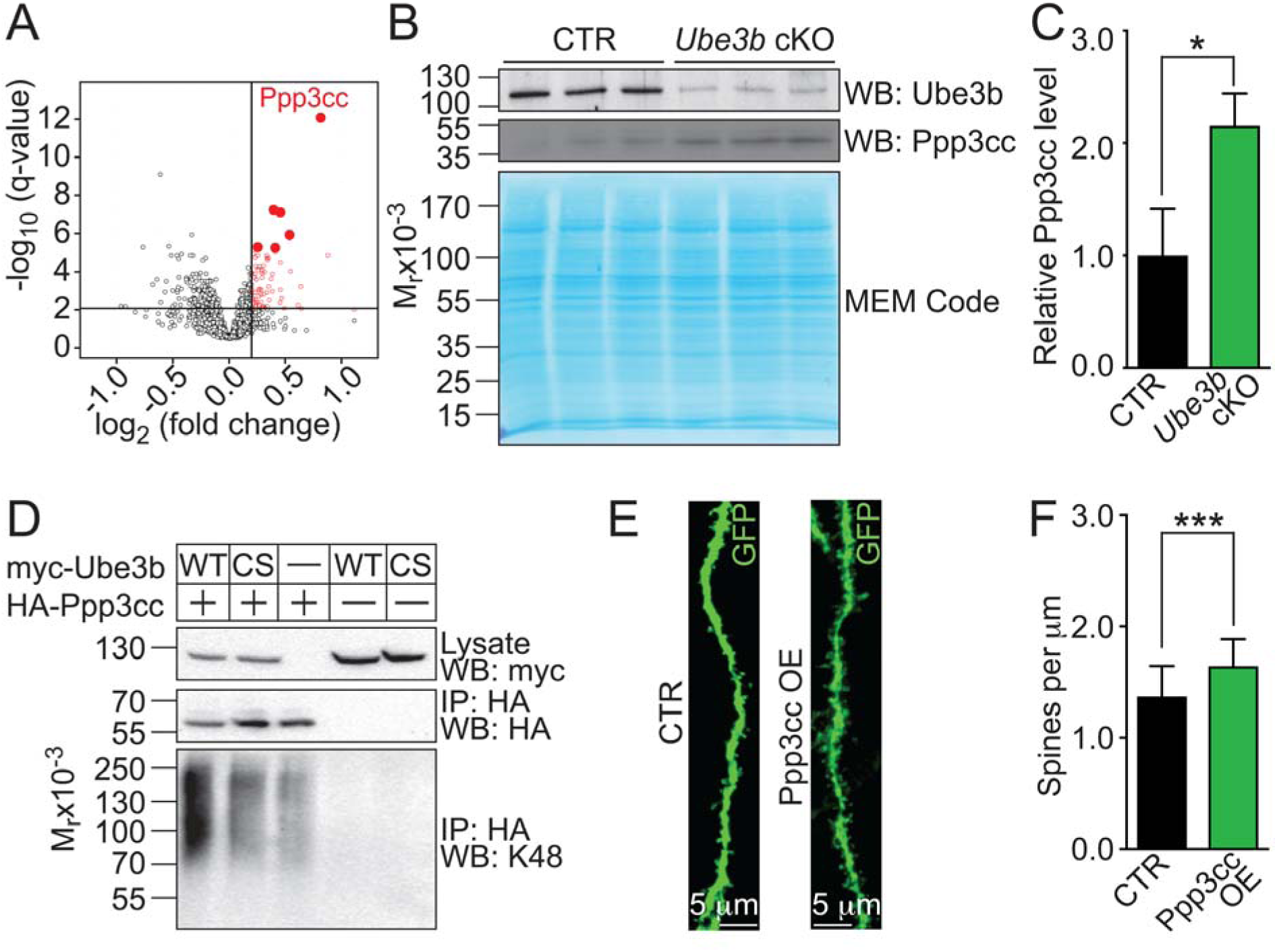
Proteomic screen identifies Ppp3cc as a Ube3b substrate. (A) Volcano plot of proteins identified in the SM3 fraction from *Ube3b* cKO cortices by quantitative proteomics. Proteins with a fold change > 1.15 and q-value < 0.01 were considered as putative Ube3b targets and are labeled in red. Top hits are indicated in red filled circles. (B) Validation of mass spectrometric quantification by Western blotting using SM3 fractions from CTR and *Ube3b* cKO with antibodies against Ube3b and Ppp3cc. MEM Code staining of nitrocellulose membrane was used as a loading control (C) Quantification of Ppp3cc levels in control and *Ube3b* cKO SM3 fractions. Protein level was normalized to MEM Code staining intensity and expressed relative to Ppp3cc level in CTR (Table S1I’). (D) Cell-based ubiquitination assay. Myc-tagged Ube3b was expressed with or without HA-tagged Ppp3cc in HEK293T cells. HA-tagged Ppp3cc was immunoprecipitated and blotted for HA (middle panel) and K48-chains of Ub (bottom panel). Note enrichment of K48-linked ubiquitination when wild-type Ube3b was co-expressed with Ppp3cc. (E) Representative images of EGFP fluorescence signals after immunostating with anti-GFP antibody from primary branches of apical dendrites from hippocampal CA1 neurons. Neurons were transfected with expression vectors for EGFP and myr-Venus alone (CTR) or EGFP, myr-Venus and HA-tagged Ppp3cc (Ppp3cc OE) by IUE at E14.5 and fixed at P21. (H) Quantification of spine density in CTR and Ppp3cc OE neurons (Table S1J’). Data are represented as averages ± S.D. For statistics on (A), see Table S2; (C) and (F), unpaired t-test; * p < 0.05.

To test if Ube3b catalyzes ubiquitination of Ppp3cc, we performed a cell-based ubiquitination assay. We expressed myc-tagged Ube3b together with HA-tagged Ppp3cc in HEK293T cells. As a negative control, we expressed catalytically inactive Ube3b C/S. As a positive control, we assayed the autoubiquitination of HA-tagged Ube3b (Fig. S6). Expression of myc-tagged Ube3b, but not of Ube3b C/S, led to an increase in K48-linked polyUb signal from immunoprecipitated HA-Ppp3cc in Western blotting (Fig. 8F). This indicates that Ube3b mediates ubiquitination of Ppp3cc and its proteasomal degradation.

### Ppp3cc overexpression phenocopies *Ube3b* cKO in spine formation

In the last set of experiments, we tested if the overexpression (OE) of recombinant Ppp3cc mimics the consequence of *Ube3b* loss in spine formation. We overexpressed recombinant Ppp3cc in wild type hippocampal neurons by IUE. Like in *Ube3b*-deficient neurons, we observed an increase in spine density in Ppp3cc OE neurons as compared to control neurons (Fig. 8E and 8F; Table S1J’). Collectively, phenocopy of *Ube3b* cKO by Ppp3cc OE supports the hypothesis that Ube3b-dependent ubiquitination and subsequent downregulation of Ppp3cc are critical for regulation of correct spine density in the developing hippocampus.

## DISCUSSION

### Dysfunction of human *UBE3B* underlies KOS, a developmental disorder with severe ID

In this work, we report that Ube3b, the murine ortholog of UBE3B is indispensable for proper neuronal development and synaptic transmission. Moreover, dorsal telencephalon-specific *Ube3b* cKO leads to alterations in circuit physiology and also in spatial and social memory. We show that Ube3b ubiquitinates Ppp3cc and thereby controls its expression level, regulating spine density in CA1 hippocampal neurons. Our study provides novel molecular insights into the cellular and molecular pathology in the KOS ID mouse model and contributes to a better understanding of the phenotypic changes in KOS patients.

### KOS protein Ube3b is a morphoregulatory Ub ligase enriched at the synapses in pyramidal neurons

*UBE3B* was discovered and cloned as one of genes up-regulated after acoustic trauma in the chick inner ear (Gong et al., 1996). Abundant Ube3b transcript in the CNS (Fig. 1A and 1B) and the developmental levels of Ube3b protein (Fig. 1C) resembled expression pattern of synaptic proteins, such as neuroligin-1 and PSD-95 (Song et al., 1999). Accordingly, we report an enrichment of Ube3b protein in the PSD fraction in a way even more pronounced than that of PSD-95 (Fig. 1D). In our *in vitro* experiments (Fig. 2 and 3), myc-Ube3b distributed diffusely in the entire primary neuron at DIV7. This is likely due to very high expression levels and/or the fact that DIV7 neurons have yet to establish functional postsynaptic densities.

We demonstrate that deletion of *Ube3b* resulted in defects in neurite development, which may contribute to the development of ID in patients (Fig. 2). This indispensable role of Ube3b was cell-autonomous and its activity-dependent, as re-expression of recombinant Ube3b, but not a catalytically inactive point mutant restored dendritic arborizations. Moreover, expression of pathogenic UBE3B mutants (i.e. G779R and R997P) in *Ube3b* cKO neurons failed to restore proper neurite branching (Fig. 3). Both mutations have been reported in homozygous configuration in patients with typical KOS. Intriguingly, both mutations localize to the HECT domain of UBE3B, indicative of detrimental effects on its catalytic function. Moreover, genetic deletion of *Ube3b* leads to alterations of spine morphology and spine density, the latter being reversible by reintroducing Ube3b to hippocampal neurons *in vivo* (Fig. 4 and S4). Our data indicate that Ube3b is an essential neuronal morphoregulatory molecule that might operate locally at synapses.

### Physiological and morphological defects of synapses in *Ube3b* cKO mice

Aberrant development and plasticity of synaptic circuits constitutes the primary cause of ID (Ramakers, 2000). Altered numbers of synaptic connections, or altered connectivity in ID mouse models are often linked to alterations of dendritic arborization and spine density/structure (Dierssen and Ramakers, 2006). Further, synaptic activity induces dynamic changes of spine shape, size, and numbers and such plasticity is strongly associated with learning (Engert and Bonhoeffer, 1999). This indicates that Ube3b-mediated changes of plasticity at the level of single neurons might lead to disrupted activity of neuronal networks which may underlie the ID-like phenotypes in *Ube3b* cKO animals.

We found increased spine density and mEPSC frequency in *Ube3b* cKO neurons, indicating more functional synapses in the absence of *Ube3b* (Fig. 5H). Additionally, increased NMDAR to AMPAR ratio in *Ube3b* cKO neurons (Fig. 5P) indicates either increased abundance of NMDARs or increased glutamate-evoked NMDAR conductance. The first possibility is supported by the enlarged spine heads in *Ube3b* KO animals (Fig. 4E and 4G), given the positive correlation of NMDAR currents and the diameter of the spine head (Noguchi et al., 2005).

Greater spine density and increased spine neck length (Fig. 4D, 4E, and 4G) are typical of immature neurons. During early synaptogenesis, the number of synapses increases and then declines in the subsequent stage of synapse elimination; a process dependent on the input from afferent neurons. Synapse elimination selects the survivor synapses to establish the neuronal network. Disturbances in synaptogenesis and/or synapse elimination are often proposed causative for ID inpatients and observed in ID-mouse models with dendritic spines with elongated necks (Comery et al., 1997). It is possible that loss of *Ube3b* in mice leads to the retention of juvenile features of dendritic spines into adulthood. Such changes might be the basis of neurological defects of KOS.

### Divergent changes in hippocampus-dependent behaviors in *Ube3b* cKO mice

Conventional *Ube3b*^-/-^ mice generated here exhibited a multitude of developmental anomalies, including difficulties in movement, failure to thrive and severe skeletal and organ malformations, also reported previously. Phenotypes in these mice seem to originate from metabolic diseases, suggesting a severe systemic dyshomeostasis (Cheon et al., 2019). In order to provide the molecular insights into the origin of ID in our animal model and in KOS patients, we studied neuronal, cell-autonomous roles of Ube3b using forebrain-specific cKO mice. *Ube3b* cKO animals did not exhibit motor phenotypes (Fig. S5C), therefore allowing to assess, among others, higher cognitive functions. We report that *Ube3b* cKO in mice disrupts spatial learning and memory while enhancing sociability and improving social memory (Fig. 7).

Spatial memory impairment is associated with morphological changes of the hippocampal CA1 region (Tsien et al., 1996). Studies on spine shapes have classically been performed in cells impregnated by rapid Golgi method or by confocal imaging of fluorescent neurons. In the current study, we report STED microscopy-based analysis of spines from ID-mouse model, demonstrating detailed morphological changes and increased density of dendritic spines in CA1 pyramidal neurons in *Ube3b* cKO animals. Our observations may indicate a straightforward and detrimental contribution of altered morphology of CA1 neurons to circuits important for spatial learning and memory.

Animal behavior is associated with the activity of specific neuronal networks resulting in oscillatory patterns of activity. Synchronized activity of neuronal circuits enables communication between different regions of the CNS and establishes a biological substrate for higher cognitive functions of the cerebral cortex. Hippocampal γ-oscillations in mice are triggered during exploratory behavior and navigation (Csicsvari et al., 2003). Increased power of γ-oscillations in *Ube3b* cKO mice may reflect aberrant activity of neuronal networks resulting in profound behavioral alterations in *Ube3b* cKO mice. The CA2 region of the hippocampus is essential for social memory and interaction in mice (Hitti and Siegelbaum, 2014). It is tempting to speculate that, *Ube3b* loss exerts detrimental effects on neuronal wiring in the CA1 region implicated in spatial learning and improves neuronal function in the CA2 subfield necessary for social memory. Our results corroborate the model that the neural substrates of spatial and social memory are distinct and can be affected independently.

Reported Ube3b-mediated loss of spatial memory and improvement in social interactions seem to be a unique feature of the KOS mouse model. Entirely opposite behavioral alterations were reported for a neuroligin-3 R451C mutant mouse, which exhibits superior spatial learning and impaired social interactions, characteristic features for autism spectrum disorders (ASDs) (Tabuchi et al., 2007). Based on these animal models, KOS seems to represent an ID syndrome with pathology different from ASDs.

### Ppp3cc as a substrate of Ube3b

We report upregulation of Ppp3cc, a γ-isoform of catalytic subunit of protein phosphatase 2B, also known as calcineurin in SM3 fractions of *Ube3b* cKO brain (Fig. 8). There are three genes encoding the catalytic subunits of calcineurin, α, β, and γ-isoform. Interestingly, calcineurin has been described as a key regulator of synaptic transmission, due to its ability to negatively regulate opening probability, current and insertion into the membrane of glutamate receptors. Calcineurin is known to regulate glutamatergic transmission by indirectly influencing the NR2B/NR2A subunit composition of NMDARs (Lin et al., 2011). Moreover, calcineurin-mediated dephosphorylation of metabotropic glutamate receptor mGluR5 enhances glutamate-evoked currents through NMDAR (Alagarsamy et al., 2005; Lin et al., 2011). The observed increase in NMDAR to AMPAR ratio in *Ube3b* cKO might be attributable to increased levels of Ppp3cc and its effects on glutamatergic transmission.

Ppp3cc is important for brain plasticity, and its dysregulation due to pathological Ca^2+^ signaling has been associated with cognitive brain disorders, such as schizophrenia (Horiuchi et al., 2007). Its Ca^2+^-dependence is mediated by calmodulin. Interestingly, Ube3b is also regulated by calmodulin through its direct Ca^2+^-independent binding to IQ motif of Ube3b (Braganza et al., 2017). Further research should elucidate Ca^2+^/calmodulin signaling and Ube3b in the aspects of brain plasticity that underlie complex brain function.

Interestingly, morpholoregulatory effects of calcineurin-mediated dephosphorylation of Growth-Associated Protein 43 (GAP43) are associated with abrogated neurite growth (Lautermilch and Spitzer, 2000). Overexpression of non-phosphorylatable GAP43 leads to spatial learning deficits in the water maze test in mice (Holahan and Routtenberg, 2008). It is tempting to speculate that loss of Ube3b leads to increased Ppp3cc levels and thereby mediates neurite branching deficits in a GAP43-dependent manner.

Ppp3cc has been described in dendritic spines (Wu et al., 2012), where it localizes to F-actin and regulates cytoskeleton stability (Halpain et al., 1998). Our data demonstrate that an increase in Ppp3cc either by *Ube3b* KO or by overexpression of recombinant Ppp3cc protein leads to an increase in spine density.

These features make Ppp3cc a promising Ube3b substrate, underlying the molecular determinants of complex phenotypes in our ID mouse model, the *Ube3b* cKO.

In conclusion, our results provide evidence for a novel indispensable regulatory machinery, whereby Ube3b-dependent ubiquitination of Ppp3cc controls formation of dendritic spines necessary for proper synchronization of the local neuronal networks. Our model demonstrated here provides molecular and cellular insights into developmental underpinnings to KOS.

## AUTHOR CONTRIBUTIONS

MCA: conceptualization, data curation, formal analysis, methodology, writing and editing of the original draft; SR, EB, MS, TS, BA, LP, SH, OJ, ED, MR, HE, KW, JR: data curation, methodology, and formal analysis; GB, RY: resources and conceptualization; VT: editing and review of the draft; HK: conceptualization, project administration, editing and review of original draft.

## ACKNOWLEDGEMENT

This work was supported by the German Research Foundation (SPP1365/KA3423/1-1 and KA3423/3-1, HK; DFG TA 303/4-2, VT), and the Russian Scientific Foundation (15-14-10021, VT), JSPS KAKENHI Grant Numbers 15K21769 (HK), The Mother and Child Health Foundation (HK), and the Fritz Thyssen Foundation (HK). We would like to acknowledge Nils Brose for his help with project administration and insightful remarks regarding the conceptualization. We thank Fritz Benseler, Klaus-Peter Hellman, Bernd Hesse-Niessen, Ivonne Thanhäuser, Dayana Warnecke, Christiane Harenberg, Maik Schlieper, Dörte Hesse, Rike Dannenberg and Denis Lajkó and animal facilities of Max Planck Institute of Experimental Medicine and of Charité University Hospital. We acknowledge Ahmed Mansouri and Judith Stegmüller for their feedback.

## SUPPLEMENTARY TABLES

Supplementary Table 1 (related to the entire paper). Exact values, n number and statistics for experiments presented in this report.

Supplementary Table 2 (related to Fig. 8). Label-free quantification of proteins in SM3 fractions from control and *Ube3b* cKO cortices.

## MATERIALS AND METHODS

### CONTACT FOR REAGENTS AND RESOURCE SHARING

Further information and requests for resources and reagents should be directed to and will be fulfilled by Hiroshi Kawabe.

#### Animals

Colonies of wild type mice (*Mus musculus*) have been maintained in the animal facilities of Max Planck Institute of Experimental Medicine (MPIem) and Charité University Hospital. *Ube3b*^f/f^ mouse line was generated in this study by using homologous recombination in ES cells (The European Conditional Mouse Mutagenesis Program, EUCOMM). Animals homozygous for the loxP alleles were viable, fertile, and born at the expected Mendelian ratio, and exhibited no overt phenotypic changes in the cage environment. To inactivate *Ube3b* in the developing cortex, we crossed *Ube3b*^f/f^ mice with the *Emx1*-Cre line (B6.129S2-Emx1^tm1(cre)Krj^/J), in which Cre recombinase is expressed from the *Emx1* gene allele (Gorski et al., 2002). *Ube3b*^f/f^, *Ube3b*^f/f^; *Emx1*-Cre^+/-^, and *Ube3b*^+/-^ (obtained by crossing the *Ube3b*^f/f^ mice with B6.FVB-Tg(EIIa-cre)^C5379Lmgd^/J, *E2A*^Cre/+^ line) (Lakso et al., 1996) were maintained in C57BL/6N background.

All experiments conducted on mice were performed in MPIem and Charité University Hospital in compliance with the guidelines for the welfare of experimental animals approved by the State Government of Lower Saxony (Niedersächsisches Landesamt für Verbraucherschutz und Lebensmittelsicherheit, LAVES), and the Max Planck Society for experiments in MPIem and by the State Office for Health and Social Affairs, Council in Berlin, Landesamt für Gesundheit und Soziales (LAGeSo) for experiments in Charité University Hospital. These guidelines are comparable to National Institute of Health Guidelines.

#### Sex and age/developmental stage of animals for *in vivo* experiments

Female and male KOS patients are both equally affected by the loss of *UBE3B*. For this reason, littermates of both sexes were randomly assigned to experimental groups during the procedure of IUE, biochemical, electrophysiological and cell biological experiments. For the behavioral analyses and quantification of cortical thickness, neuron number and inflammatory response, we analyzed male mice.

Analysis of the cortical thickness, neuron number in the cortex and inflammatory response was performed using 9 week old animals. Cortical layering and corpus callosum was assessed using P0 mouse brains. Dendritic spines in *Ube3b* cKO, *Ube3b* KO and Ppp3cc OE neurons were quantified after IUE at E14.5 on P21 brains. The rescue of dendritic spine density using myc-Ube3b expression in Cre-transfected neurons was performed at P10. Finally, the behavior analyses were carried out using animals at 8 weeks old at the beginning of testing.

#### Cell lines and primary cultures

##### HEK293FT and HEK293T cell line

HEK293FT were used for production of lentiviruses and for transfections (Gibco; R700–07); HEK293T were used for transfections (the details are specified in he figure legends or in the main text. HEK293FT, or HEK293T cells were maintained in 6 cm Petri dishes (Corning) in 10 mL HEK cell medium at 37°C in the presence of 5% carbon dioxide in HERA-cell240 (Heraeus) incubator. For passaging, semi-confluent cells were washed with PBS, incubated with 1 mL of 0.05% trypsin solution (Gibco, Life Technologies) for 1 minute at 37°C and the reaction was stopped by addition of 9 mL of the fresh medium. Sex of HEK293FT/293T cells is female.

###### HEK cell medium

500 mL DMEM, 50 mL FBS, 5 mL GlutaMAX, 5 mL penicillin/streptomycin (100X).

##### Murine primary neurons

Primary hippocampal cultures used in this paper were prepared from P0 mouse brains, exactly as described (Ambrozkiewicz et al., 2018).

##### Mouse Embryonic Fibroblasts (MEFs)

Neomycin-resistant MEFs (Cell Biolabs; CBA-311) as feeder cells for embryonic stem (ES) cell culture. MEFs were maintained in 15 cm Petri dishes coated with gelatin (0.1% gelatin in ddH_2_O, Sigma-Aldrich) at 37°C in 25 mL MEF medium in the presence of 5% carbon dioxide in HERA-cell240 (Heraeus) incubator. At confluency, medium was removed, cells were washed with 37°C-warm PBS, and incubated with 10 mL MEF medium supplemented with 100 μL of mitomycin C (Sigma-Aldrich) per plate for 2.5 hours at 37°C to mitotically inactivate MEFs. Then, medium was removed, cell washed with PBS, trypsinized for 5 minutes with 0.05% trypsin-EDTA solution (Gibco, Life Technologies) at 37°C, resuspended with fresh MEF medium, and centrifuged at 1 krpm for 5 minutes at room temperature. Inactivated MEF were then resuspended, Freezing medium was added dropwise to the suspension, and cells were frozen in Cryo Freezing Container (Nalgene) filled with isopropanol at −80°C, and seeded prior to demand.

###### MEF medium

500 mL KnockOut-DMEM (Gibco, Life Technologies), 75 mL FBS, 6 mL non essential amino acids (Gibco, Life Technologies), 6 mL 200 mM L-glutamine (Gibco, Life Technologies), 6 mL β-mercaptoethanol (Sigma-Aldrich), 3 mL penicillin/streptomycin, 6 mL G418 (Gibco, Life Technologies).

###### Freezing Medium

22 mL MEF medium, 10 ml FBS, 8 mL dimethyl sulfoxide (DMSO, Sigma-Aldrich).

##### Embryonic Stem Cells (ESCs)

For generation of Ube3b knockout mice, mutant JM8.F6 ES cells were purchased from EUCOMM Consortium. The L1L2_gt0 cassette was inserted to the genome at position 114389925 (UCSC Genome Bioinformatics) of Chromosome 5 upstream of exon 7 of Ube3b gene. The cassette was composed of an FRT-flanked lacZ/neomycin sequence followed by a 5’ loxP site. An additional loxP site is inserted 3’ downstream of exon 7 at position 114390710. ES cells were thawn prior to blastocyst injection to minimize the passage number. Cells were seeded on a layer of previously prepared and thawn inactivated MEF, and maintained in ES cell medium at 37°C in the presence of 5% carbon dioxide in HERA-cell240 (Heraeus) incubator. Before splitting, cells were supplied with fresh medium for at least 3 hours. Passaging was carried out with 0.25% trypsin-EDTA (Gibco, Life Technologies), and the reaction was terminated with FBS. Prior to blastocyst injection, MEFs were separated from ES cells by plating cell suspension on fresh gelatinized Petri dish for 30 minutes at 37°C allowing MEFs to adhere to the bottom of the dish. Transfer of injected blastocysts to pseudo-pregnant foster female mice was performed in MPIem Animal House Facility.

###### ES cell medium

500 mL KnockOut-DMEM, 95 mL FBS, 6 mL non essential amino acids, 6 mL 200 mM L-glutamine, 6 mL β-mercaptoethanol, 3 mL penicillin/streptomycin, 6 mL G418, 1000 U Leukemia Inhibitory Factor (Chemicon/Millipore).

### METHOD DETAILS

#### Generation of *Ube3b^f^*^/f^ mouse line

ESCs were validated with PCRs and Southern blotting (Fig. S1A to S1D) and injected into blastocyst (Thomas and Capecchi, 1987). Mice derived from such ES cells were crossed with flip recombinase expressing mouse line (B6;SJL-Tg(ACTFLPe)9205Dym/J; Jackson Laboratory) to delete the lacZ/neomycin cassette. *Ube3b*^f/+^ mice were crossed to establish *Ube3b*^f/f^ mouse line.

#### Mutant mouse genotyping

Genotyping for all mutant mouse lines was performed in DNA Core Facility (MPIem, Göttingen) by PCR using oligonucleotide primers listed below (wt, wild type allele; Cre, knock-in allele with Cre; ko, knockout allele; floxed, allele with loxP site).

*Emx1*^Cre/+^: 5’-GAAACAGCCCCCTGCTGTCC -3’; 5’- AGCCAGCCCATTCTCTTGTCCCTC - 3’; 5’-CATCGCCACGAAGCAGGCCAACGG -3’; wt – 234 bp, Cre – 267 bp, *Ube3b*^f/f^: 5’-TCGGGTGTTTACTTGGATAACTCT -3’; 5’-TGTGCTTTGGTTCCTTATCTGTC-3’; 5’-CCACAACGGGTTCTTCTGTTAG -3’; wt – 197 bp, floxed – 345 bp.

The genotyping PCRs were carried out using the following parameters:

Step 1: 96°C for 3 minutes,

Step 2: 94°C for 30 seconds,

Step 3: 64°C for 1 minute,

Step 4: 72°C for 1 minute (32 cycles from Step2 to 4),

Step 5: 72°C for 7 minutes.

Multiplex PCR to detect 3’loxP site (Fig. S1C) was performed with these parameters using oligonucleotide primers listed for *Ube3b*^f/f^ genotyping. Amplification of long genomic DNA sequences for validation of correct recombination of targeting vector to genomic DNA, (Fig. S1B) was performed using Pfu-AD polymerase in Phusion-HF buffer (NEB) in the presence of 1 M betaine (Sigma-Aldrich), and oligonucleotides 5’-GTATCTTATCATGTCTGGATCCGGGGG-3’ and 5’-ATGCTGGCAGACTTTGCACTCTTTACTCTC -3’ with the following parameters:

Step 1: 99°C for 3 minutes,

Step 2: 99°C for 30 seconds,

Step 3: 60°C for 30 seconds,

Step 4: 72°C for 90 seconds (30 cycles from Step 2 to 4)

Step 5: 72°C for 2 minutes.

#### Vectors

##### pCR2.1-SB-3’Ube3b

To generate Southern blot probe for validation of the homologous recombination of Ube3b targeting vector, primers 5’-TTAGTGTGGCTTTTCAGCCTTAA-3’ and 5’- TGGAGCCGTTAGGTCATTTCA-3’ were used for PCR using DNA isolated from wild type ES cells as a template. Amplified DNA was subcloned into pCR2.1-TOPO.

##### pCR2.1-HECT-Ube3b

To generate a template for in situ hybridization probe, primers 5’-CGGATGTTGGAGGACGGCTA-3’ and 5’-CTTGATGATGGAACGGAAGCC-3’ were used in PCR to amplify cDNA encoding for Ube3b HECT domain using P0 cortex cDNA library as a template. Amplified DNA was subcloned into pCR2.1-TOPO.

##### pCR2.1-Ube3b (WT, full length)

cDNA NM_054093.2 (Entrez) encoding full length Ube3b (1070 aa) was amplified with primers 5’-GAGGAATTCACCATGTTCACTGTATCTCAGACCTCCAGAGC-3’ and 5’-CTCCTCGAGTTATTAGGAGAGCTCAAAGCCCGTGTT-3’, introducing 5’EcoRI and 3’XhoI restriction sites using pYX-Asc-Ube3b (BioCat) as a template. Resulting DNA was then subcloned into pCR2.1-TOPO.

##### pCAG-myc-Ube3b

pCR2.1-Ube3b was digested with EcoRI and XhoI, and ligated with pCAG-myc-1 opened with the same restriction enzymes.

##### pCR2.1-Ube3b C1038S

Primers 5’-GGCGTCTGCCCACATCTTCCACTAGCTTCAACCTGCTTAA -3’ and 5’-\TTAAGCAGGTTGAAGCTAGTGGAAGATGTGGGCAGACGCC-3’ were used for site-directed mutagenesis to generate pCR2.1-Ube3b C/S introducing point C1038S mutation into pCR2.1-Ube3b WT.

##### pCAG-myc-Ube3b C1038S

pCR2.1-Ube3b C1038S was digested with EcoRI and XhoI, and ligated to pCAG-myc-1 opened with the same restriction enzymes.

##### pcDNA3-HA-hUBE3B, pcDNA3-HA-hUBE3B G779R, pcDNA3-HA-hUBE3B R997P

Human Ube3b cDNA was amplified in PCR reaction using primers 5’-AAAAAAGATCTATGTTCACCCTGTCTCAGACC-3’ and 5’-AAAAATCTAGACTAGGAGAGTTCAAAGCCCG-3’ and cloned into pcDNA3 vector using BglII and XbaI restriction sites and T4 DNA ligase, which was then further used to introduce single aminoacid substitution to generate point mutants G779R and R997P using the following primer pairs, respectively: 5’-GCTCTTCGAGTTTGTGAGGAAGATGCTGGGGAA-3’ and 5’-TTCCCCAGCATCTTCCTCACAAACTCGAAGAGC-3’ (for G779R); 5’-CCATTCTCCATCCCCTGCGTGGAGGTG-3’ and 5’-CACCTCCACGCAGGGGATGGAGAATGG-3’ (for R997P).

##### pCAG-HA-Ppp3cc

cDNA NM_001304991.1 (Entrez) encoding full length Ppp3cc was amplified with primers 5’- AGATTACGCTATCTGTACAGAATTCACCATGTCCGTGAGGCGC-3’ and 5’-GGCCGCTAGCCCGGGTACCGAATTCTTACAGGGCTTTCTTTCCATGGTC-3’ and inserted into pCAG-HA vector using NEB Builder (NEB).

Oligonucleotides used in this study were synthetized by DNA Core Facility of Max Planck Institute of experimental Medicine. Other plasmids used in this study: pCAG (gift from Jun-ichi Miyazaki), pRaichu205X (gift from Michiyuki Matsuda), pCX::myrVenus (gift from Anna-Katerina Hadjantonakis), and pYX-Asc-Ube3b (BioCat)

#### Antibodies

Primary antibodies used in this publication are as following (dilution for Western blotting, WB; for immunocytochemistry, ICC; for immunohistochemistry, IHC; for immunoprecipitation, IP): rabbit anti-Ube3b, 339003, SySy (WB, 1:1000); mouse anti-actin, AC40, Sigma-Aldrich (WB, 1:2000); mouse anti-PSD-95, NeuroMab, K28/43 (WB: 1:1000); rabbit anti-RabGDI, 130011, SySy (1:1000); rabbit anti-GFP, 598, MBL (IHC and ICC, 1:2000); goat anti-GFP, 600-101-215M, Rockland (IHC and ICC, 1:1000); mouse anti-myc, 9E10, Sigma-Aldrich (ICC, 1:500); mouse anti-myc, 9B11, Cell Signaling Technologies (IP, 1:1000); rabbit anti-myc, 9B10, Cell Signaling Technologies (WB, 1:1000); rabbit anti-HA, C29F4, Cell Signaling Technologies (WB, 1:1000); mouse anti-HA, Covance (ICC, 1:1000); mouse anti-NeuN, Millipore (IHC, 1:400); rabbit anti-Satb2, self made (Ambrozkiewicz et al., 2017) (IHC, 1:1000); rat anti-CTIP2, 25B6, abcam (IHC, 1:500); rat anti-L1, Millipore (IHC, 1:2000); rabbit anti-Cux1, M222, SCBT (IHC, 1:500); rabbit anti-Ppp3cc, PA5-15583, Thermo (WB, 1:500); mouse anti-ubiquitin, P4D1, SCBT (WB, 1:1000); rabbit anti-K48, D9D5, Cell Signaling Technology (WB, 1:1000); rabbit anti-K63, D7A11, Cell Signaling Technology (WB, 1:500). Staining of cellular nuclei was performed using DAPI or Draq5 (Biostatus Limited). HRP-coupled secondary antibodies used in this study were from BioRad: goat anti-mouse (1721011) and anti-rabbit (1706515) IgG HRP. For quantitative Western blotting with Oddyssey system, IR-Dye coupled antibodies were used (Li-COR): anti-mouse IgG IR-Dye-680RD (925-68070), anti-mouse IgG IR-Dye-800CW (925-32210), anti-rabbit IgG IR-Dye-680RD (925-68071), anti-rabbit IgG IR-Dye-800CW (925-32212). Fluorophore-coupled secondary antibodies were from Thermo Fisher Scientific and Jackson Immunoresearch.

#### Southern blotting

Restriction digest of 10 μg of genomic DNA isolated from ES cells was held using 10U of SapI restriction enzyme (NEB) in 37°C for 16 hours. Next, digested genomic DNA was resolved in 0.7% agarose gel in 500 mL TAE buffer supplied with 40 μL of 1% ethidium bromide (Carl Roth) at 30V for 16-20 hours. After gel documentation in UV (Intas), agarose gel was incubated in 0.25 M HCl (Merck) for 10 minutes, and subsequently in 0.4 M NaOH (Merck) for 10 minutes. Upward capillary transfer of DNA onto a nylon membrane (Hybond N+, GE Healthcare) was performed in 0.4 M NaOH for 16-20 hours. Next, the membrane was incubated in 2XSSC until neutralized, and air dried on chromatography paper (Whatman). DNA was further UV-crosslinked with energy density of 1.00J/cm^2^ (FluoLink, Biometra), and the membranes were pre-hybridized in Rapid Hyb-Buffer (GE Healthcare, Amersham) 1 hour at 65°C. Radioactive probes were synthetized with Prime-It II Random Primer Labeling Kit (Agilent) using 25 ng of EcoRI-digested pCR2.1-SB-3’Ube3b and α-^32^P-labeled dCTP (Perkin-Elmer) following manufacturer’s protocol. After termination of probe synthesis, the sample was diluted with MB-H_2_O to final volume of 100 μL, passed through chromatography columns to remove unincorporated dCTP with Bio-Spin size exclusion column (Bio-Rad). 25 μL of the eluate was diluted with 5 mL of pre-warmed (65°C) hybridization buffer, and incubated with DNA crosslinked on the nylon membrane. Hybridization was held at 65°C for 2 hours, followed by extensive washing with 2XSSC buffer with 0.1% SDS (sodium dodecyl sulfate, Gerbu) for 15 minutes in room temperature, and subsequently with 1XSSC buffer with 0.1% SDS for 7 minutes at 65°C. Radioactivity of the membrane was then monitored with Geiger-Müller counter, and the washing was continued until the scintillator reported approximately 100 counts. Membrane was then exposed to photographic film (Kodak) for 16-20 hours at −80°C, and developed.

##### TAE buffer

40 mM Tris-base (Sigma-Aldrich), 20 mM acetic acid (Sigma-Aldrich), 1mM EDTA (Sigma-Aldrich).

##### 1XSSC buffer

150 mM NaCl (Merck), 15 mM sodium citrate (Merck), pH 7.0.

#### Subcellular fractionation of mouse brains

Subcellular fractionation of the brain tissue was performed in sucrose gradients. All described procedures were performed at 0-4°C. Immediately after decapitation, mouse cortex was isolated in Solution A. Cortices from one mouse were homogenized in 3 mL of Solution A supplemented with 0.2 mM PMSF, 1 μg/mL aprotinin, and 0.5 μg/mL leupeptin using Teflon-glass homogenizer with 20 strokes at 1200 rpm (Potter S, Braun). Homogenate (H) was fractionated by centrifugation at 82 500 *g* for 2 hours in discontinuous sucrose density gradient, comprising of 0.85 M, 1.0 M, 1.2 M sucrose solutions. The solution on top of the first interface was collected as supernatant (S), the interface between 0.32 and 0.8 M as the P2A fraction (myelin fraction), the one between 0.8 and 1.0 M as the P2B fraction (ER/Golgi fraction), the one between 1.0 and 1.2 M as the P2C fraction (synaptosomes), and the pellet as the P2D (mitochondria-enriched fraction) fraction. The P2C fraction was centrifuged at 100 *kg* for 20 minutes, and the pellet was resuspended with the hypo-osmotic buffer (6 mM Tris-HCl, pH 8.0) followed by incubation for 45 minutes to disrupt the synaptosomes by the hypo-osmotic shock. The sample was further centrifuged at 32.8 *kg* for 20 minutes to separate the synaptic cytoplasm, and crude synaptic vesicles (SC/CSV) fraction in the supernatant from the crude synaptic membrane (CSM) fraction in the pellet. The CSM fraction was further resuspended with 6 mM Tris-HCl pH 8.0, 0.32 M sucrose, and 0.5% Triton X-100 (Roche), incubated for 15 minutes, and centrifuged at 32.8 *kg* for 20 minutes. The pellet was harvested as postsynaptic density fraction (PSD), and the supernatant – Triton X-100 extract (TX100-E).

For the purpose of comparative mass spectrometry, CSM fractions were subjected to additional centrifugation in discontinuous sucrose gradient as described above. The interface between 1.0 and 1.2 M sucrose solutions was collected as the synaptic plasma membrane (SM3) fraction.

##### Solution A

0.32 M sucrose (Merck), 1 mM NaHCO_3_ (Merck).

#### HEK293FT cell transfection and Ubiquitylation assay

Transfection of HEK293FT was performed at 80-90% confluency using Lipofectamine2000 (Invitrogen, # 11668027) following manufacturer’s instructions with 1:1 DNA/Lipofectamine2000 ratio (μg/μL). Cells were maintained in 6 cm Petri dishes and transfected with 12 μg DNA per condition with 1:1 ratio between plasmids encoding myc-Ube3b variants and HA-Ppp3cc. 24 hours post-transfection, after removal of the medium, cells were gently washed with PBS supplemented with 20 mM N-ethylmaleimid (NEM) and dishes with cells were flash frozen in liquid nitrogen. Next, cells were lysed in 150 μL of Lysis Buffer A and the homogenate incubated in 65°C for 20 minutes. After that, the homogenate was diluted with 1.2 mL of Lysis Buffer B and subjected to centrifugation at 10 000 rpm for 10 minutes. Further, the supernatant was incubated with 25 μL HA-beads (Roche, # 11 815 016 001) at 4°C overnight. After subsequent five washing steps with Lysis Buffer B, the bead volume was adjusted to 50 μL using Lysis Buffer B and further to 75 μL using Lämmli buffer.

##### Lysis Buffer A

50 mM Tris-Cl pH 7.5, 300 mM NaCl, 1% SDS, 1 μg/μL Aprotinin, 0.5 μg/μL Leupeptin, 0.2 mM PMSF, 10 mM NEM.

##### Lysis Buffer B

50 mM Tris-Cl pH 7.5, 300 mM NaCl, 1% Triton X-100, 1 μg/μL Aprotinin, 0.5 μg/μL Leupeptin, 0.2 mM PMSF, 10 mM NEM.

### Image acquisition, analysis, and statistics

#### Quantitative Western blotting

Quantification of protein levels was performed with ImageStudioLite (Li-Cor) software using scans of films after exposure to ECL chemiluminescence or data acquired by Odyssey cLX. The bands representing respective proteins were manually outlined, and the signal intensity was measured. Signal represents the sum of the individual pixel intensity values for a selected shape subtracted by the average intensity value of the pixels in the background, and the surface area of the region of interest. Protein level was expressed relative to the signal for β-actin detected in respective lane or to the total protein as measured by the integral of MEM Code Staining intensity, which was measured with Tracing tool of ImageJ software (Schindelin et al., 2012). For statistical analysis of difference between two groups, t-test was used, and p-value of less than 0.05 was considered as significant.

#### Spine morphometrics with STED nanoscopy

Spine morphology was quantified with ImageJ software using images acquired with STED microscope as described before (Willig et al., 2014). Spines were manually counted using Cell Counter plug-in of ImageJ software, and classified into morphology classed: stubby, filopodia, thin, mushroom, and bifurcated using morphological criteria described before (Harris et al., 1992). Spine length was measured as the distance from spine emergence on the dendrite until the head tip. The head size was measured as the diameter of spine head. For statistical analyses, t-test for comparison of two groups, and one-way ANOVA with Bonferroni post-hoc test for analysis of spine classification were conducted.

#### Quantification of axon length and dendritic tree complexity

Axon length was measured using confocal images of neurons acquired by Leica Sp2 with 20X objective and ImageJ software. Number of primary branches were counted manually using Cell Counter plug-in. To measure the complexity of dendritic arbor, images of primary hippocampal neurons were acquired with AxioImager Z.1 (Carl Zeiss), using 40X objective and water immersion. Next, we applied Sholl analysis with 7.5 μm interval between Sholl circles with median span type on tresholded and binarized images using ImageJ. For quantification of dendritic tree complexity, we quantified a total number of crossings with Sholl circles made by neurites of an individual neuron. For statistical analyzes, t-test was used to compare two independent samples, and one way ANOVA with Bonferroni post-hoc tests for comparison of more than two groups.

#### Confocal imaging of immunostained brain sections and analysis

Cortical thickness of *Ube3b*^f/f^ and *Ube3b*^f/f^;*Emx1*-Cre^+/-^ was measured as the distance from the pia to the end of layer VI on coronal cross section. Tresholded binarized images after Watershed segmentation algorithm using ImageJ were used for quantification of number of neurons. NeuN-positive punctae more than 50 μm^2^ in size were counted as somata. For statistical analysis to compare two groups, t-test was employed.

To acquire representative images of primary branches of CA1 apical dendrites of neurons expressing EGFP and myrVenus, Leica Sp5 microscope with 40X objective with confocal scanning Zoom 2.5 was used. Quantitative imaging was performed using ImageJ measure tool on manually selected somata.

### Electrophysiology

#### Autaptic neurons

Briefly, autaptic cultured neurons (9-14DIV) were whole-cell voltage–clamped at −70 mV with the amplifier EPSC10 (HEKA) under the control of the Patchmaster 2 program (HEKA) (Ripamonti et al., 2017). All acquired traces were analyzed using AxoGraph X (AxoGraph Scientific). Recordings were performed with a patch-pipette solution and the extracellular solution (BASE+Ca^2+^+Mg^2+^). EPSCs was evoked by depolarizing the cell from −70 to 0mV at a frequency of 0.2 Hz. The ready releasable pool (RRP) size was measured after 6 s application of M hypertonic sucrose solution. The vesicular release probability, Pvr, was calculated as a ratio of the charge transferred during an action potential induced-EPSC divided by the charge during sucrose application. Short-term plasticity was measured by recording EPSC during 10 Hz stimulation. Miniature EPSC (mEPSC) was recorded in the presence of 300 nM tetrodotoxin (TTX, Tocris Bioscience). Surface expression of glutamate, and GABA receptors was measured by application of 100 μM glutamic acid (Sigma) or 3 μM GABA (Sigma), respectively. AMPA and NMDA ratio were calculated diving the charge transfer before and after bath application of a Mg^2+^-free, Ca^2+^-and-glycine-containing solution.

##### Patch-pipette solution (all reagents from Sigma-Aldrich)

138/16.8 mM potassium gluconate/Hepes pH 7.4 at 310 mOsm, 10 mM NaCl, 1 mM MgCl_2_, 0.25 mM EGTA, 4 mM ATP-Mg^2+^, 0.3 mM GTP-Na^+^.

##### BASE+Ca^2+^+Mg^2+^

140 mM NaCl, 2.4 mM KCl, 10 mM Hepes, 10 mM glucose, 4 mM CaCl_2_, and 4 mM MgCl_2_ at 320mOsml/liter, pH 7.3.

#### **γ**-Oscillation recording

γ-oscillations were recorded in *Ube3b*^f/f^ and *Ube3b*^f/f^;*Emx1*-Cre^+/-^ mice at P15-23. Transverse hippocampal sections (300 µm thickness) were prepared from age-matched littermates as described previously (Ripamonti et al., 2017) using a Leica VT1200S vibratome. Brain isolation and slice preparation were performed in sucrose-based slicing solution at 4°C in the presence of carbogen. After sectioning, slices were placed in a holding chamber containing artificial cerebrospinal fluid (ACSF) and let recover for 15 minutes before recording. γ-oscillations were recorded in CA3 field of hippocampus, where the peak power of kainate-induced γ-oscillations is the highest. Interface recording chamber (BSCBU Base Unit with the BSC-HT Haas Top, Harvard Apparatus) was used during recordings. Slices were steadily perfused with ACSF, and 33°C temperature was maintained. Extracellular recording electrode filled with ACSF was inserted into CA3 pyramidal cell layer, and extracellular field potential were recorded for more than 20 minutes before the addition of 50 nM or 100 nM kainate. Baseline (20 minutes), and oscillatory activity (30 minutes) were measured using a 700B amplifier (Axon Instruments, Molecular Devices), low pass Bessel filtered at 3 kHz, and digitized by the Digidata 1440A data acquisition system (Axon Instruments, Molecular Devices). All traces were analyzed using Axograph X software (AxoGraph Scientific). Power spectra were calculated upon Fourier transforms of 10-minutes epochs (last 10 minutes of each recording) of recorded field activity. The baseline power spectrum was subtracted from the power spectrum obtained during kainate application. The dominant frequency within 25-45 Hz was used to quantify the peak frequency. The power was calculated as the area under the respective peak in the power spectrum.

##### Sucrose-based slicing solution

230 mM sucrose, 26 mM NaHCO_3_, 2 mM KCl, 1 mM KH_2_PO_4_, 2 mM MgCl_2_, 10 mM D-glucose, 0.5 mM CaCl_2_.

##### ACSF

120 mM NaCl, 26 mM NaHCO_3_, 1 mM KH_2_PO_4_, 2 mM KCl, 1 mM MgCl_2_, 10 mM D-glucose, 2 mM CaCl_2_.

Other procedures used in this paper such as *In situ* hybridization, Immunocytochemistry, Immunohistochemistry, *In utero* electroporation, Protein concentration measurements, Sodium dodecyl sulfate polyacrylamide gel electrophoresis (SDS-PAGE), Calcium phosphate transfection of hippocampal primary neurons, Lentivirus production and infection of primary hippocampal cells, Perfusion, Cryosectioning, Western blotting, and Comparative Label Free Mass Spectrometry were performed exactly as thoroughly described in our recent publication (Ambrozkiewicz et al., 2018).

### Behavioral testing

#### Animals and housing conditions

All behavioral experiments were approved by the local animal care and use committee in accordance with the German animal protection law. For behavioral testing, mice were housed in groups of 5 in standard cages, with food and water available ad libitum. Animals were kept under a 12h light-dark cycle with lights on at 7:00am and an ambient temperature of 20-22°C.

In the course of behavioral experiments, we noticed that when *Ube3b*^f/f^; *Emx1*-Cre^+/-^ males were crossed to *Ube3b*^f/f^ females, the offspring were positive for a full *Ube3b* knockout allele (“rec” in Fig. S1), likely due to germline leakage of Cre expression in this *Emx1*-Cre^+/-^ driver line. For this reason, we only present the data for males born from *Ube3b*^f/f^; *Emx1*-Cre^+/-^ females crossed to *Ube3b*^f/f^ males. Offspring are all negative for the “rec” allele in the genomic DNA samples prepared from the tail tissue.

#### Behavioral phenotyping

All behavioral experiments were conducted by an experimenter who was unaware of the genotype of the mice. Experiments were performed during the light phase of the day (between 8:00am and 5:00pm). Male *Ube3b* cKO mice (N=11) and their control littermates (N=8) were tested in an extensive behavioral test battery. The behavioral test battery was performed as previously described (Dere et al., 2014; Netrakanti et al., 2015) and included the following tests: Visual cliff test, hearing assessment, olfaction, pain perception, tape test, LABORAS, open-field, rotarod, elevated plus-maze, marble burying, pre-pulse inhibition, water-maze, nesting, social interaction in pairs, and sociability and social memory in the tripartite chamber. The age of mice at the beginning of testing was 8 weeks.

### Sensory function

#### Vision (Visual cliff test)

The visual cliff test consisted of a perspex box (70×35×30 cm) that had a transparent floor. It was placed on the edge of a laboratory bench, so that 50% of its base was positioned on the bench (’ground’ side), while 50% of the base protruded over the edge of the bench, suspended 1m above the floor (’air’ side). The mouse was placed in the middle of the ground side and the time spent on both sides of the box over a period of 5 minutes was measured using a video-tracking system (Viewer2, Biobserve, Germany).

#### Hearing assessment based on acoustic startle reflex

In order to assess the integrity of the sensorimotor reflex system that mediates the startle reflex to a sudden loud noise, we determined a detailed tone intensity-startle response curve. The startle reaction to an acoustic stimulus (pulse) is a short-latency reflex that is mediated by an oligosynaptic neural circuit that includes the lower brainstem, spinal and cranial motor neurons, and the cerebellum.

The startle response was measured in a startle box. The startle reflex induces a movement of a force-sensitive platform, which was recorded over a period of 100 ms, beginning with the onset of the pulse. An experimental session consisted of a 2 minutes habituation period to a 65dB background white noise, followed by a baseline recording for 1 minutes. After baseline recording, stimuli of different intensity and a fixed duration of 40 msec were presented. Stimulus intensity was varied between 65dB and 120dB, such that 19 intensities (in steps of 3dB) from this range were used. Every single stimulus intensity was presented 10 times in a pseudorandom order with an inter-stimulus interval of 8-22s. The amplitude of the startle response (expressed in arbitrary units) was defined as the difference between the maximum force detected during the recording window and the force measured immediately before the stimulus onset. For each mouse, the amplitudes of the startle responses for a given stimulus intensity were averaged.

#### Olfaction (Buried food finding test)

The mice were first habituated to the test cage (29.5×18.5×13 cm) for 3 consecutive days with 2 daily trials of 20 minutes duration each. On days 4 to 6 the mice had been food deprived for approximately 22h prior to the 2 daily habituation trials. The mice received a piece of chocolate cookie (1.6g) during each habituation trial and additionally 3-5 cookies in its home cage after testing. After the second habituation trial the mouse had access to food in its home cage for 1h. On day 7 the mice had to locate a piece of chocolate cookie that was hidden beneath approximately 1.5 cm below fresh bedding close to the wall at one end of the cage. Each mouse was placed into the right corner at the opposite end of the cage, and the time it needed to locate the cookie and to start burying to recover it was measured with a cut-off time of 3 minutes. The mouse was removed from the test cage after the cookie had been discovered and before it was consumed. As a control for possible motivational deficits, a visible test trial was performed after the hidden food test. Again the latency to locate the cookie was measured with a cut-off of 3 minutes.

#### Pain perception (Hot-plate test)

The hot plate test was used as a measure of pain perception. Mice were placed on a metal plate (UgoBasile Srl, Comerio, Italy) that was preheated up to 55°C. The latency to show hind-paw raising, licking or jumping was recorded. Immediately after showing such indications of discomfort, the mice were removed from the platform. A cut-off time of 40 sec ensured that no mouse was injured.

#### Somatosensory perception (Tape-test)

The tape or adhesive removal test measures the integrity of somatosensory perception in mice. In preparation of the tape test the forepaws of the mouse were cleaned with water and thereafter thoroughly dried with a paper towel. Thereafter a small adhesive tape (dimensions 0.3 x 0.4 cm) was placed randomly on top of either the left or right forepaw of the mouse. The time needed to sense the tape and to remove the adhesive from the paw was measured. If a mouse was not able to remove the adhesive after 120 sec the tape was removed by the experimenter and a score of 120 was noted.

### Motor function, spontaneous and exploratory activity

#### Motor coordination and balancing (Rotarod)

This test measures motor coordination, balancing and motor learning. The rotarod consists of a horizontal drum (ENV-577M, Med Associates Inc. Georgia, Vermont, USA) that rotates along its vertical axis. Over a period of 5 minutes, the drum was accelerated from 4 to 40 rpm. Each mouse was placed individually on the drum and the latency until it slid off the drum was recorded using a trip-switch. 24h later the mouse was re-tested to measure motor learning.

#### Spontaneous home-cage activity (LABORAS)

The LABORAS home-cage behavior monitoring system (Metris b.v., Hoofddorp, The Netherlands) consisted of a triangular shaped sensor platform (Carbon Fiber Plate 1000×700×700×30 mm), positioned on 2 orthogonally placed force transducers (Single Point Load Cells) and a 3rd fixed point attached to a heavy bottom plate (Corian Plate 980×695×695×48 mm). The apparatus was mounted on 3 spikes, which were adjustable in height and absorb external vibrations. Mice were housed in transparent polycarbonate cages (Makrolon type II cage, 22×16×14 cm) with a floor covered with wood-chip bedding. The cage was placed directly onto the sensor platform, with the upper part of the cage (including the top, food hopper and drinking bottle) suspended in a height-adjustable frame separate from the sensing platform. Mechanical vibrations caused by the movement of the animal were transformed by force transducers into electrical signals, amplified to a fixed signal range, filtered to eliminate noise, digitized and then stored on a computer. The computer then processed the stored data using several signal analysis techniques to classify the signals into specific behavioral categories including locomotion, rearing, movement, circling, grooming, climbing, scratching and eating. The behavior that dominates at a certain time point is scored by the LABORAS software. For each behavioral category, the time spent performing this particular behavior (counted in seconds), as well as its frequency, was measured.

#### Exploratory activity (Open-field)

The mice were placed into the center of a gray circular Perspex arena (diameter 120 cm; height of wall 25 cm). Each mouse was observed over a period of 7 minutes. The time spend in the peripheral, intermediate and center zone of the apparatus and the total distance travelled (mm), as well as the mean running velocity (mm/s) was measured using an automated tracking software (Viewer2, Biobserve, Germany).

### Psychiatric disease-related tests

#### Anxiety (Elevated plus-maze)

The apparatus was made of gray Perspex with central platform (5×5 cm), 2 open and 2 walled arms (30×5×15 cm each). Illumination density at the central platform was 135lx. The mouse was placed on the central platform and was allowed to explore the apparatus for 5 minutes. The time spent and distance travelled (mm) on the walled and open arms as well as the mean running velocity (mm/s) was measured using an automated tracking software (Viewer2, Biobserve, Germany).

#### Impulsivity (Marble burying)

Impulsivity and stereotypic behaviors were measured with the marble burying test. The mouse was introduced into a cage (34.5×56.5×18 cm) that contained standard bedding (5 cm fill height). 24 glass marbles were placed on top of the bedding. The marbles were arranged on top of the betting in 6 rows with 4 marbles per row. The distance between 2 marbles within a row was 4 cm. Trial duration was 30 minutes. The number of marbles buried was scored.

#### Sensory-motor gating (Pre-pulse inhibition)

Impairments in sensory-motor gating indicative for a neural network disturbance that is characteristic for patients suffering from schizophrenia were measured with the pre-pulse inhibition test. Mice were placed in small metal cages (82×40×40 mm) to restrict major movements and exploratory behavior. The cages were equipped with a movable platform floor attached to a sensor that recorded vertical movements of the floor. The cages were placed in 4 sound attenuating cabinets (TSE Systems, Bad Homburg, Germany). Startle reflexes were evoked by acoustic stimuli delivered by a loudspeaker that was suspended above the cage and connected to an acoustic generator. The startle reaction to an acoustic stimulus that induces a movement of a force-sensitive platform was recorded over a period of 260 msec beginning with the onset of the pulse. An experimental session consisted of a 2 minutes habituation to a 65dB background white noise (continuous throughout the session), followed by a baseline recording for 1 minutes at background noise. After baseline recording, 6 pulse-alone trials using startle stimuli of 120 dB intensity and 40 msec duration were applied to decrease the influence of within-session habituation. These data were not included in the 120dB/40 msec analysis of the pre-pulse inhibition. For tests of pre-pulse-inhibition, the startle pulse was applied either alone or preceded by a pre-pulse stimulus of 70-, 75-, or 80dB intensity and 20 msec duration. A delay of 100 msec with background noise was interposed between the presentation of the pre-pulse and pulse stimulus. The trials were presented in a pseudorandom order with a variable inter-stimulus interval ranging from 8-22 sec. The amplitude of the startle response (expressed in arbitrary units) was defined as the difference between the maximum force detected during the recording window and the force measured immediately before the stimulus onset. For each animal, the amplitudes were averaged separately for the 2 types of trials (i.e. stimulus alone or stimulus preceded by a pre-pulse). Pre-pulse inhibition was calculated as the percentage of the startle response using the following formula: % Pre-pulse inhibition=100-[(startle amplitude after pre-pulse)/(startle amplitude after pulse only)×100].

Startle response to 120dB: The response of the mouse to the startle stimulus of 120dB alone, without any pre-pulse, as well as the response without stimulus presented was also recorded.

### Spatial learning and memory

#### Morris water-maze

Hippocampus-dependent spatial learning and memory was assessed with the water maze test. A large circular tank (diameter 1.2 m and depth 0.68 m) was filled with opaque water to a depth of m. An escape platform (10×10 cm) was submerged 1 cm below the surface. The swim trajectory of the mouse was monitored by a computer and the video-tracking system VIEWER 2 (Biobserve, Germany). The escape latency, swim speed, and path length was recorded for each mouse. During the first 2 days, mice were trained to swim to a visible platform (visible platform task) that was marked with a 15 cm high black flag and placed pseudo-randomly in different locations across trials (non-spatial training). The extra-maze cues were hidden during these trials. After 2 days of visible platform training, hidden platform training (spatial training) was performed. For 8 days, mice were trained to find a hidden platform (i.e. the flag was removed) that was located at the center of one of the four quadrants of the pool. The location of the platform was fixed throughout testing. Mice had to navigate using extra-maze cues that were placed on the walls of the testing room. Every day, mice went through 4 trials with an inter-trial interval of 5 minutes. The mice were placed into the pool facing the side wall randomly at 1 of 4 start locations and allowed to swim until they found the platform or for a maximum of 90 sec. Any mouse that failed to find the platform within 90 sec was guided to the platform. The animal then remained on the platform for 20 sec before being removed from the pool. The next day after completion of the hidden platform training, a spatial probe trial was conducted. The platform was removed from the pool, and mice were allowed to swim freely for 90 sec.

The preference for the former platform zone was expressed as percent of time spent in the target zone relative to the total time spend in the pool (90 sec). Mice were also compared regarding the number of entries into the target quadrant.

### Social behavior

#### Nesting

The mice were single-housed 1h before lights were turned off. The cage contained bedding material and nesting towels. After 2 nights of habituation, nesting towels were replaced by nestlets (pressed cotton squares weighing ∼3 g). Nest building was assessed on the next morning. The remainder/leftover of the nesting material was weighted and the proficiency of nest building was rated using a scale that ranged from 1-5 with lower scores indicating poor nest building behavior.

#### Social interaction in pairs

Every mouse was first individually habituated to the testing cage (30×30×30 cm) for 10 minutes over 2 consecutive days. 3 pairs of unfamiliar mice of the same genotype were then placed into the testing cage for 10 minutes. The time the animals spent in close contact was recorded by a trained observer who was unaware of the genotype of the mice.

#### Sociability and social memory (Tripartite chamber)

Social preference and memory was tested in the tripartite chamber. The apparatus consisted of a rectangular box that was divided into 3 chambers (40×20×22 cm). The dividers were made from transparent Plexiglas and had rectangular entries (35×220 mm). The floor of the box was covered with bedding that was exchanged between trials. The test mouse was introduced into the middle chamber, with the entries to the other 2 chambers closed, and allowed to acclimatize for 5 minutes. Thereafter, a small wire cage (140×75×60 mm) containing an unfamiliar male C57BL/6N mouse of the same age and weight (stranger 1) was placed in one outer chamber. An empty wire cage was positioned in the other outer chamber. The location (outer left or right chamber) of stranger 1 was alternated between trials. After unblocking the entries to the outer chambers, the test mouse was allowed to freely move between chambers for 10 minutes. Time spent in and number of entries into each chamber were recorded by a video tracking system (Viewer2, Biobserve GmbH, Germany). Each mouse received a 2nd and 3rd trial. The 2nd trial was identical to the 1st trial except that the stranger mouse was placed into the wire cage in the other outer chamber in order to control for a possible side bias. On the 3rd trial, the test mouse was presented with the familiar stranger 1 and an unfamiliar stranger 2. Sociability and social memory indexes were calculated as follows: Sociability index = (time investigating stranger)/(time investigating stranger+time investigating empty cage)x100; Memory index = (time investigating unfamiliar mouse)/(time investigating unfamiliar+time investigating familiar mouse)x100.

#### Statistical analysis

Between-group comparisons were made by either one-way analysis of variance (ANOVA) with repeated measures or t-test for independent samples. Within-group comparisons were made via t-tests for dependent samples. Within-group tests of chance level performance using ratio or percentage calculations were performed via single group t-tests against a chance level of either 0.25 or 0.5 when indicated. Mann Whitney U, Wilcoxon tests were used if the normality assumption was violated (as assessed by the Kolmogorov-Smirnov test). All statistics were performed using SPSS v.17 (San Diego, USA) or Prism Graph Pad software. Data presented in the figures and text are expressed as mean ± S. E. M.; p-values <0.05 were considered significant.

---

**Supplementary Figure 1.**
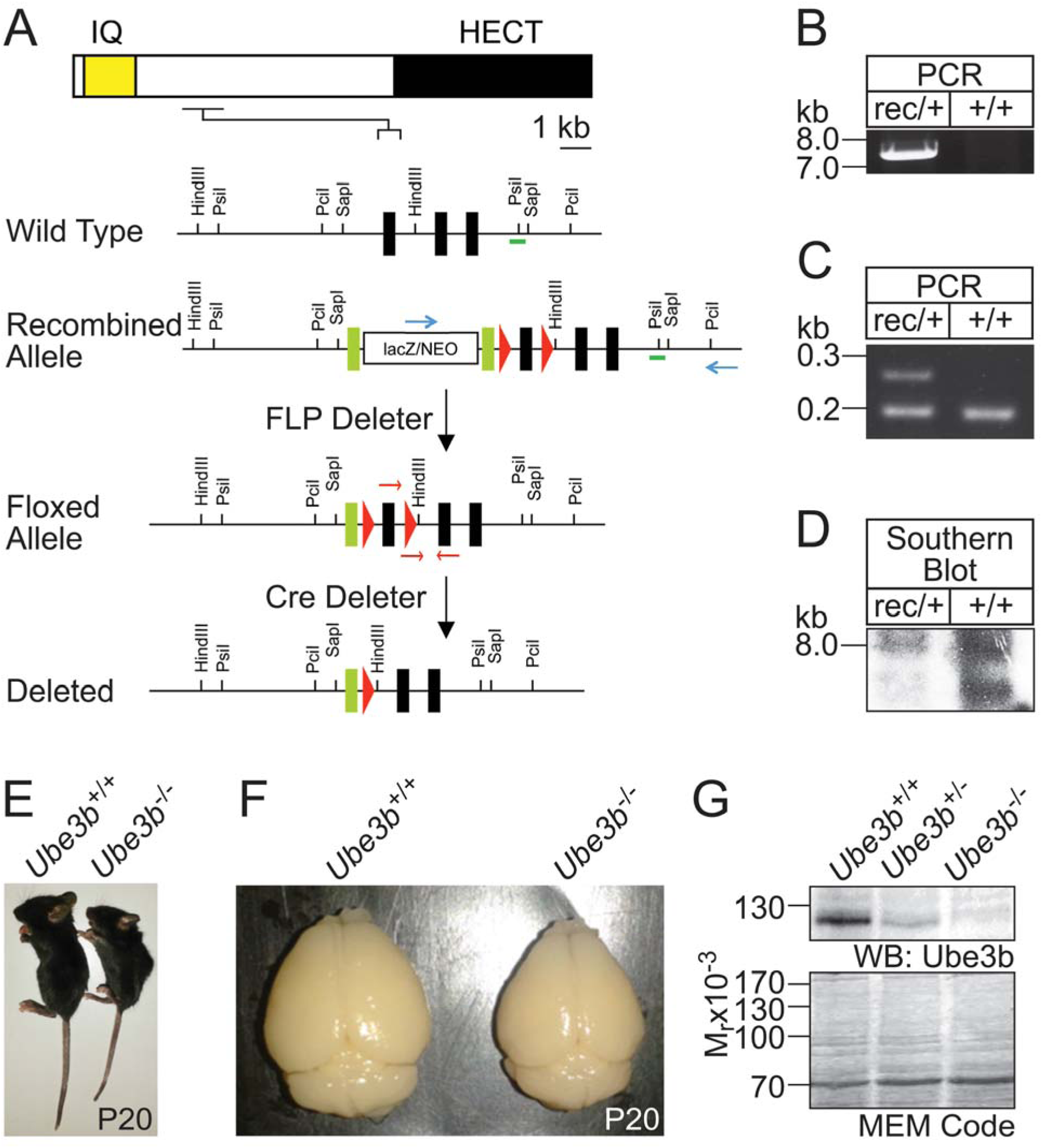
*Ube3b* knockout in mice results in growth retardation, failure to thrive and an overall smaller brain. Related to the entire paper. (A) Gene targeting strategy of murine *Ube3b*. The domain structure of Ube3b, wild type, and targeted *Ube3b* alleles are depicted. Exons, loxP sites, and FLP-recombinase target sites are symbolized as black rectangles, red triangles, and green rectangles, respectively. Exon 7 of *Ube3b* is flanked by two loxP sites. Mice positive for the recombined allele were crossed with FLP deleter animals to remove the lacZ/neomycin selection cassette, and further crossed to Cre-driver lines, leading to removal of exon 7 conditionally or conventionally. (B, C, D) Validation of *Ube3b* gene targeting. +/+ wild type, rec/+ indicates heterozygous recombined mutant. (B) The result of long-range PCR using primers depicted as blue arrows in A. Band of the predicted size was observed only using genomic DNA purified from rec/+ mutant as a template. (C) The result of multiplex PCR using primers depicted as pink arrows in A. PCR using genomic DNA from targeted ES cells yields two bands (left lane), and PCR using wild type ES cells genomic DNA as a template results in one band. (D) Southern blotting analysis of genomic DNA purified from targeted ES cells. Genomic DNA isolated from control and ES cells carrying ‘Recombined’ allele of *Ube3b* was digested with SapI enzyme. The probe is indicated as a green line in A. The band at 8.0 kb represents the wild type allele, and the band at 6.6 kb represents the mutant allele. (E) Gross morphology of *Ube3b*^+/+^ (left) and *Ube3b*^-/-^ (right) mice at 20 days post birth. (F) Brains of *Ube3b*^+/+^ (left) and *Ube3b*^-/-^ (right) mice isolated 20 days after birth. (G) Results of Western blotting using P7 brain lysates from *Ube3b*^+/+^ (left), *Ube3b*^+/-^ (middle), and *Ube3b*^-/-^ (right) animals with an antibody raised against HECT domain of Ube3b. MEM Code staining (lower panel) shows comparable amounts of total protein in all lanes.

**Supplementary Figure 2.**
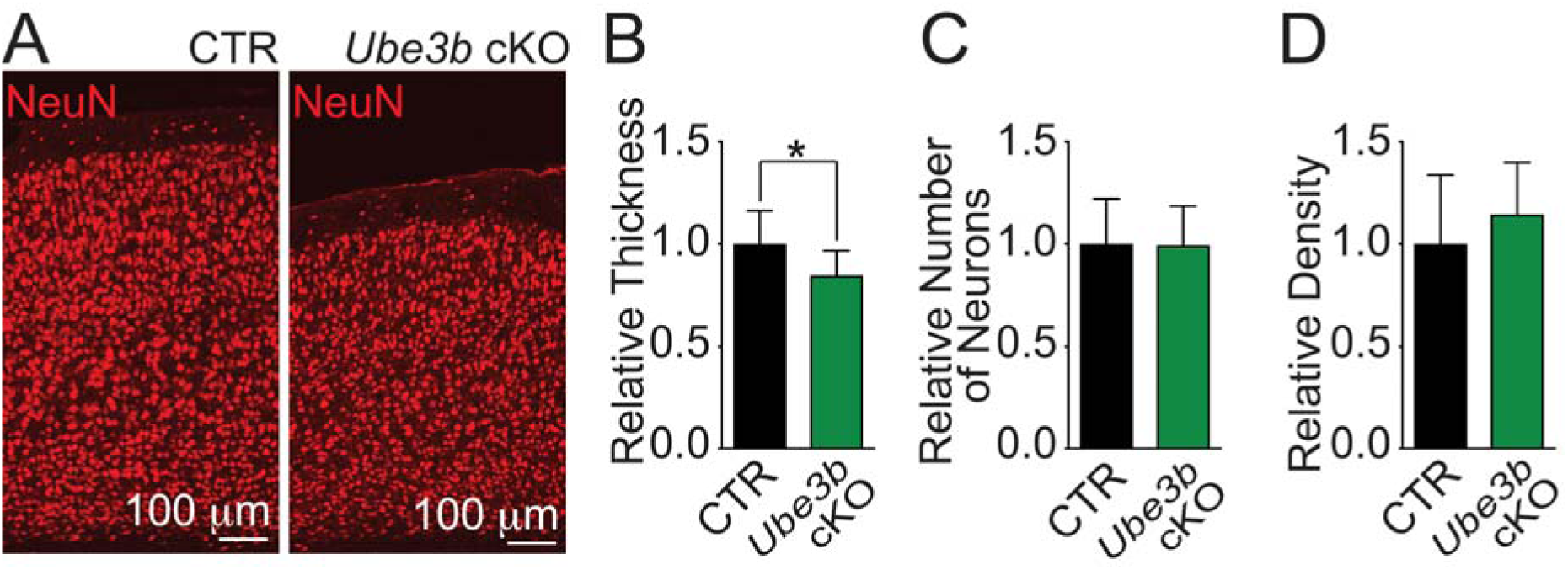
Brain-specific *Ube3b* conditional knockout mice show morphological defects in the cortex. Related to Fig. 3. (A) Reduced thickness of the cortex in *Ube3b*^f/f^; *Emx1*-Cre^+/-^ (*Ube3b* cKO) mice. Representative images of immunostaining for neuronal nuclear marker, NeuN, using coronal sections of somatosensory cortex. (B) Quantification of the cortical thickness from pial surface to the bottom edge of cortical layer VI in control and *Ube3b* cKO mice. Values are expressed relative to thickness of control cortex (Table S1F). (C) Quantification of the number of neurons in CTR and *Ube3b* cKO cortices. The number of NeuN-positive nuclei in a cortical region of 560 μm width was counted and expressed relative to the mean number in *Ube3b*^f/f^ cortex (Table S1G). (D) Quantification of neuronal density in CTR and *Ube3b* cKO cortices quantified as a number of NeuN-positive nuclei over the cortex area and expressed relative to control, *Ube3b*^f/f^ (Table S1H). Bar diagrams in (B), (C), and (D) represent average ± S.D. For statistics on (B-D), D’Agostino and Pearson omnibus normality test and Mann Whitney test; * p < 0.05.

**Supplementary Figure 3.**
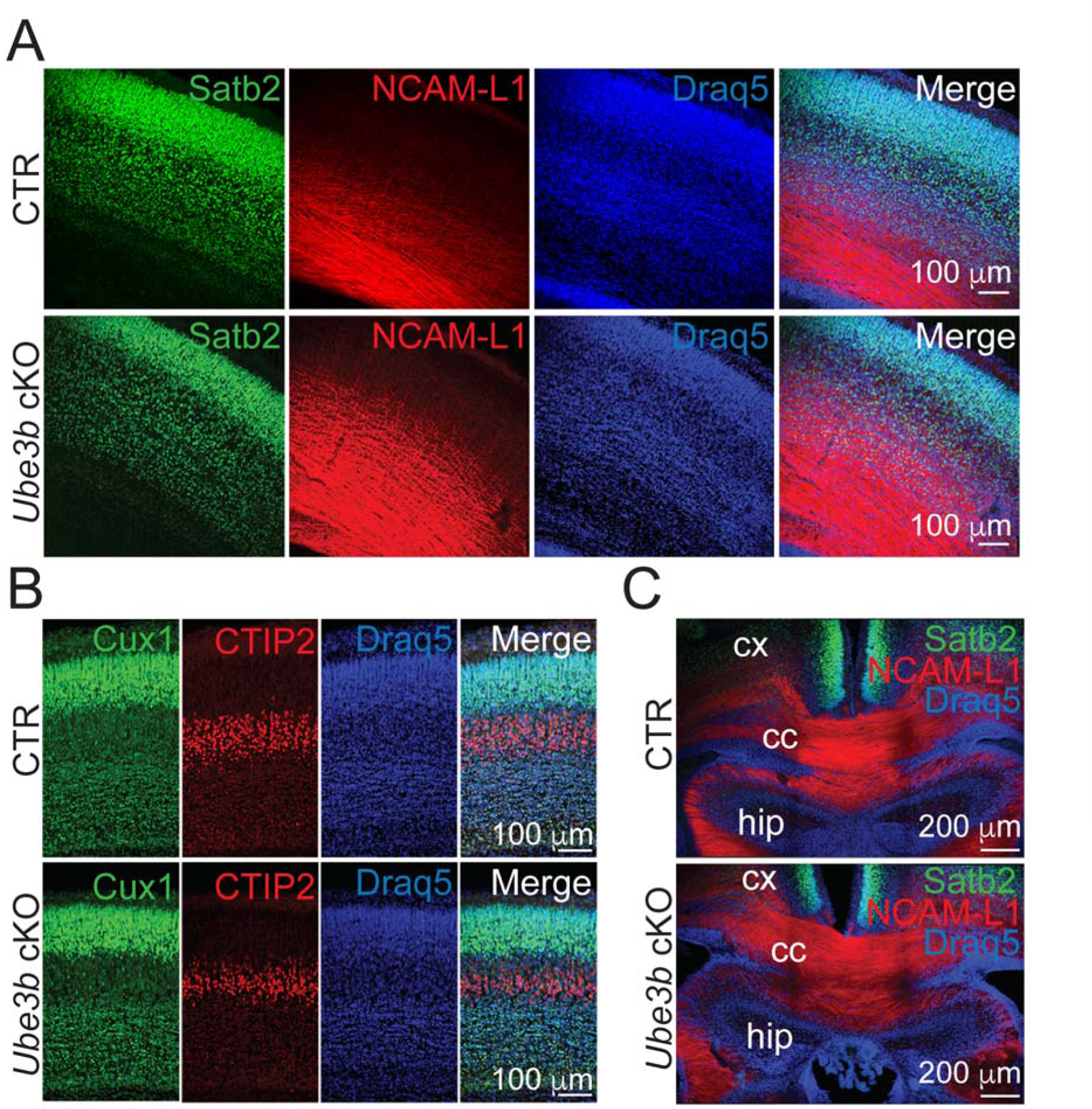
Brain-specific *Ube3b* conditional knockout mice show intact cortical layering and *corpus callosum*. Related to the entire paper. (A, B) Representative images of immunostaining with indicated antibodies. Coronal sections through somatosensory cortex of *Ube3b*^f/f^ (CTR) and *Ube3b*^f/f^; *Emx1*-Cre^+/-^ (*Ube3b* cKO) mouse were immunolabeled for Satb2 and NCAM-L1 (A), Cux1 and CTIP2 (B). (C) Coronal sections immunolabeled against Satb2, NCAM-L1 and Draq5 were also imaged at the level of the *corpus callosum*. Draq5 labels cell nuclei.

**Supplementary Figure 4.**
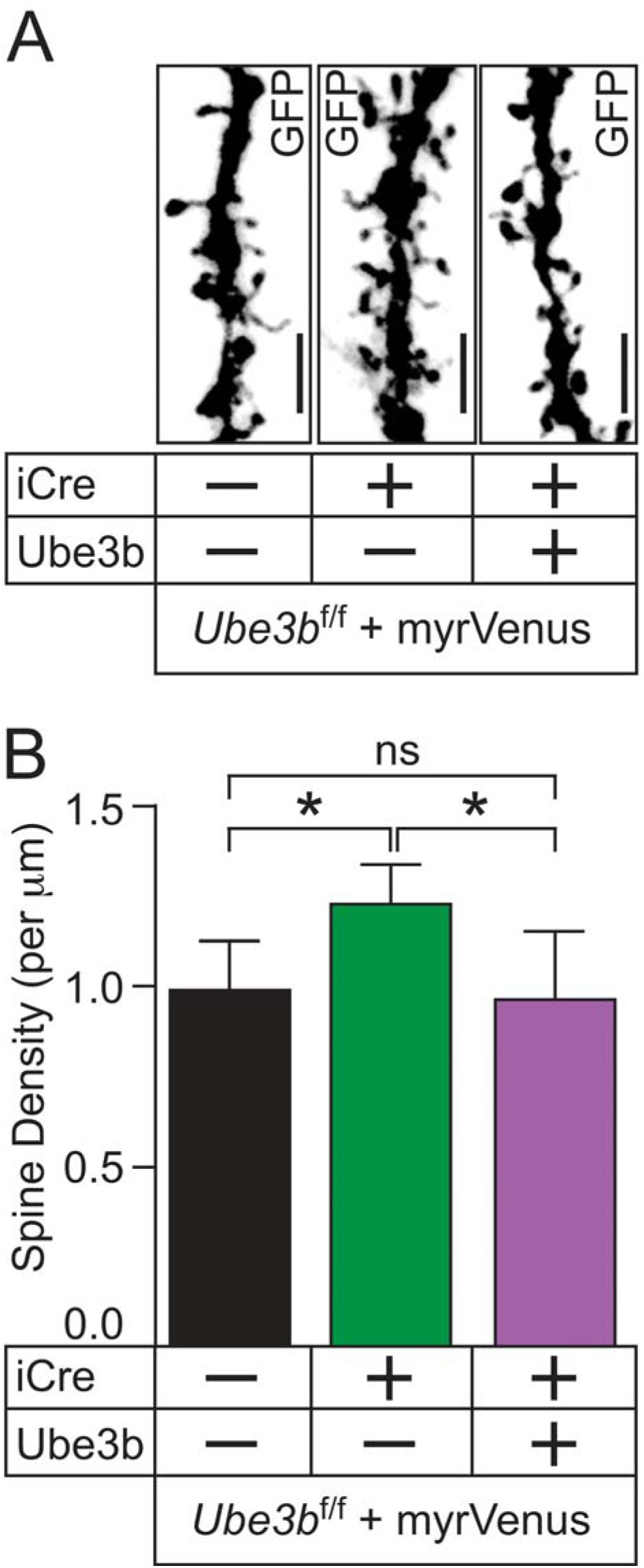
Ube3b regulates spine density in a cell-autonomous manner *in vivo*. Related to Fig. 4. (A) Representative examples immunostaining with anti-GFP antibody from primary apical branches of CA1 pyramidal neurons in *Ube3b*^f/f^ mice after confocal imaging. Anti-GFP antibody is reactive with Venus protein. Hippocampal progenitors were *in utero* electroporated at E14.5 with plasmids encoding for myr-Venus, myr-Venus and iCre, or myr-Venus, iCre and myc-Ube3b. Brains were fixed at P10 for immunostaining with the anti-GFP antibody. Scale bars, 2 µm. (B) Quantification of spine densities. Counted were numbers of spines per μm dendrite in CA1 hippocampal neurons (Table S1O). Results on bar graphs are represented as averages ± S.D. For statistics, one-way ANOVA with Bonferroni post-hoc test; * p < 0.05.

**Supplementary Figure 5.**
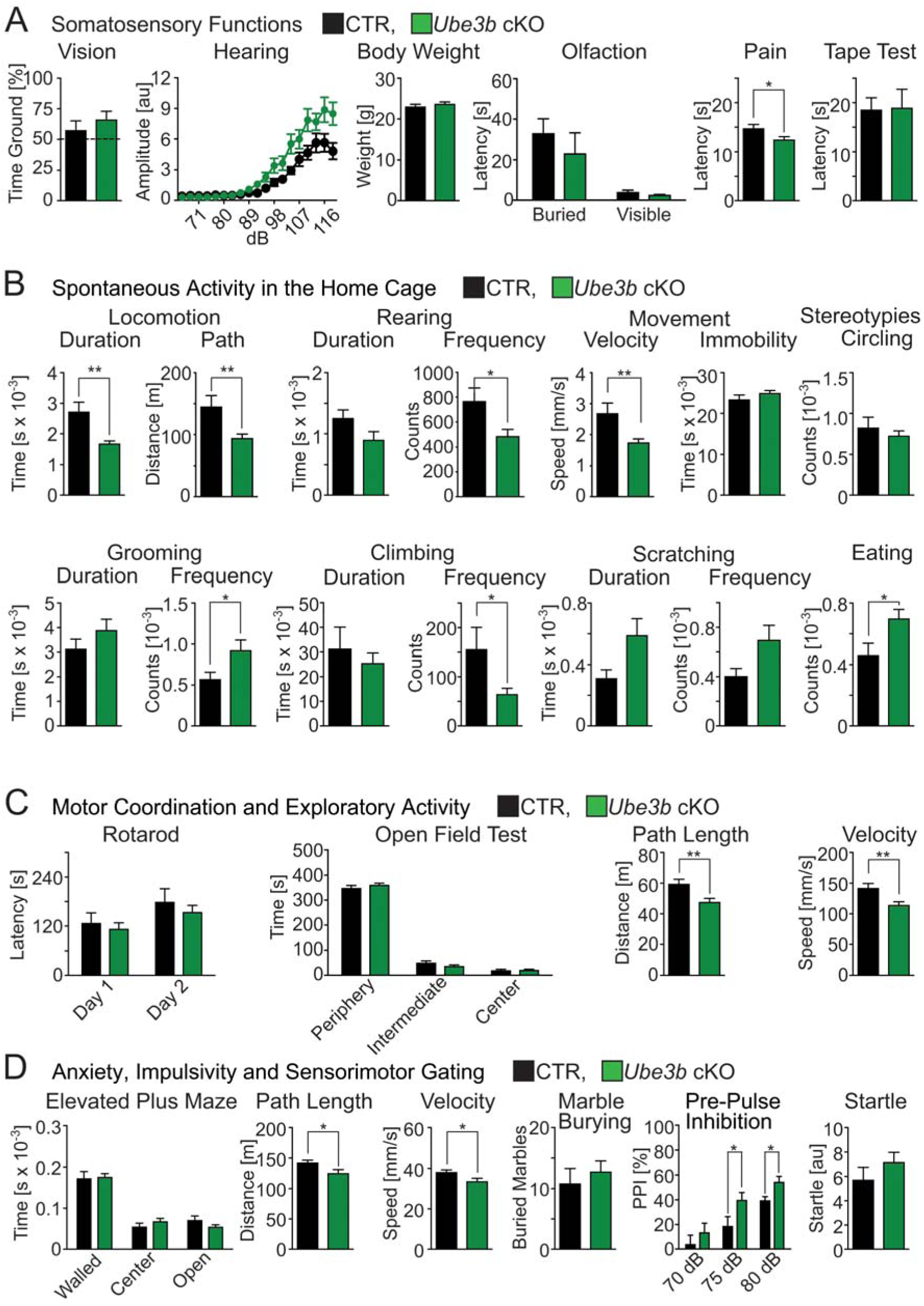
*Ube3b* cKO male mice show facilitated processing of intensive, high-arousal sensory stimuli, and behavioral switch towards self-directed actions. Related to Fig. 7. (A) Sensory functions. Quantification of vision (% time spent on the ground side of the apparatus; dashed line represents performance at chance level), hearing (startle amplitude), body weight, olfaction (latency to find the cookie), pain sensitivity (latency to show signs of discomfort), and the tape test (latency to remove the adhesive from the paw) in CTR and *Ube3b* cKO mice. (B) Home-cage spontaneous activity in the LABORAS-test. (C) Quantification of motor coordination and exploratory activity using rotarod and open-field test. (D) Quantification of anxiety, impulsivity and sensory-motor gating using elevated plus-maze test, marble burying test, pre-pulse inhibition test and startle response. For detailed information on exact experimental values and statistics, see also Table S1B’. All described tests were conducted with males. Results on graphs represent average ± S.E.M. For statistics on (A, Hearing), Repeated Measures ANOVA; otherwise, unpaired t-test; ** 0.001 < p < 0.01; * 0.01 < p < 0.05.

**Supplementary Figure 6.**
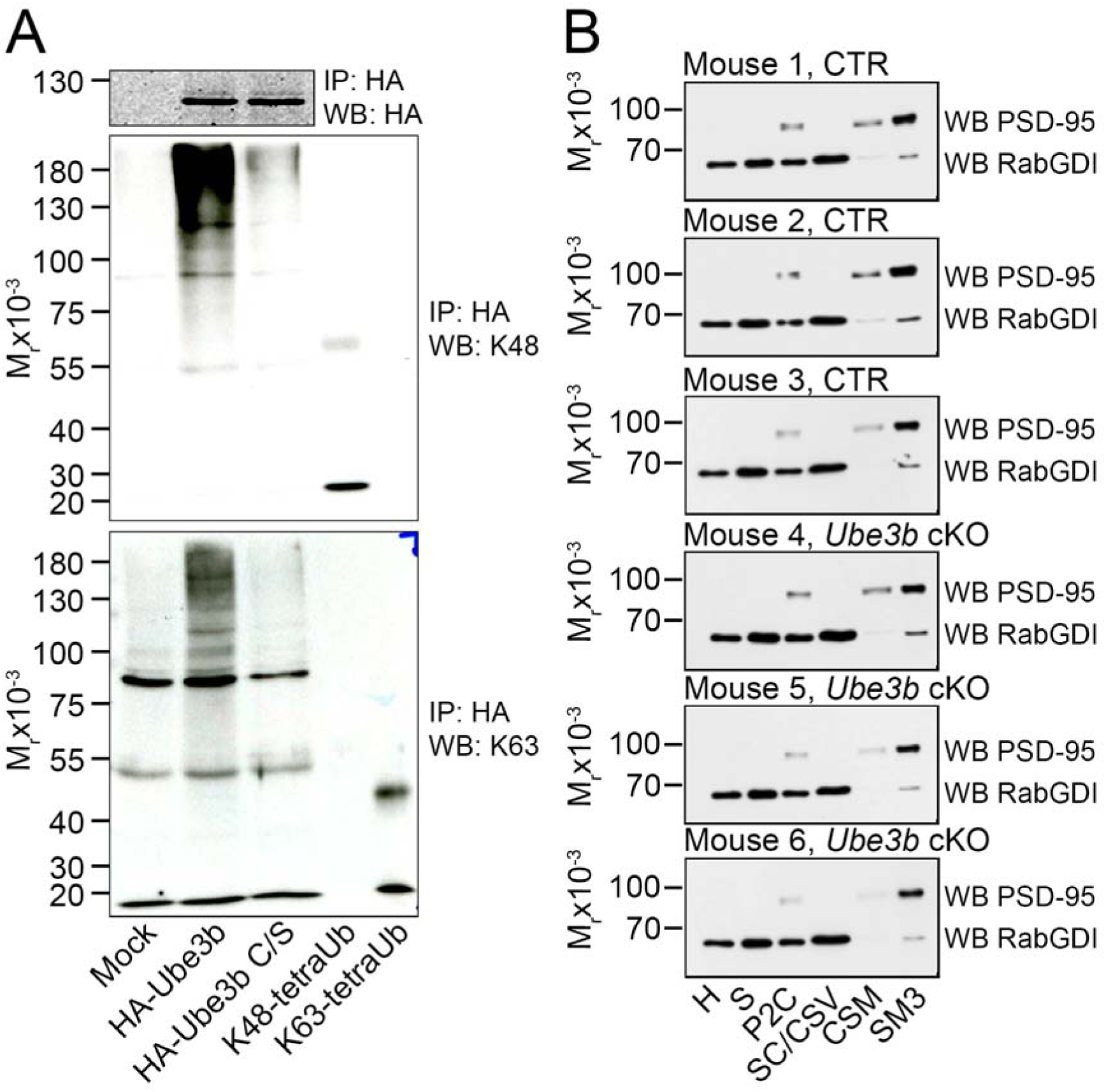
Ube3b assembles K48-linked and K63-linked Ub-chains. Related to Fig. 8. (A) HEK293 FT cells were transfected with HA-tagged wild type or catalytically inactive mutant (C/S) of Ube3b. HA-tagged Ube3b was immunoprecipitated with an anti-HA antibody, followed by Western blotting using anti-HA, anti-K48-polyUb, and anti-K63-polyUb antibodies. To compare titers of two anti-Ub antibodies, equal amounts of K48-linked and K63-linked tetra Ubs were loaded on the same SDS-PAGE gels. Note that Ube3b is able to form both K48- and K63-linked polyUb chains (second lanes in the middle and the bottom panels). (B) Western blotting validation of subcellular fractionation of CTR (Mouse 1 – 3) and *Ube3b* cKO (Mouse 4 - 6) brains using antibodies against PSD-95, and RabGDI. Homogenate (H), supernatant (S), synaptosomes (P2C), synaptic cytoplasm, crude synaptic vesicles (SC/CSV), crude synaptic membrane (CSM), synaptic membrane fraction (SM3).

